# A Strain of *Streptococcus mitis* Inhibits Biofilm Formation of Caries Pathogens via Abundant Hydrogen Peroxide Production

**DOI:** 10.1101/2024.08.06.606862

**Authors:** Isabella Williams, Jacob S. Tuckerman, Daniel I. Peters, Madisen Bangs, Emily Williams, Iris J. Shin, Justin R. Kaspar

## Abstract

Commensal oral streptococci that colonize supragingival biofilms deploy mechanisms to combat competitors within their niche. Here, we determined that *Streptococcus mitis* more effectively inhibited biofilm formation of *Streptococcus mutans* within a seven species panel. This phenotype was common amongst all assayed isolates of *S. mutans*, but was specific to a single strain of *S. mitis*, ATCC 49456. The growth inhibitory factor was not effectively carried in spent supernatants of *S. mitis*. However, we documented ATCC 49456 to accumulate 4-5 times more hydrogen peroxide (H_2_O_2_) than other species tested, and 5-18 times more than other *S. mitis* strains assayed. The *S. mutans* biofilm formation inhibitory phenotype was reduced when grown in media containing catalase or with a *S. mitis* mutant of pyruvate oxidase (*spxB*; *pox*), confirming that SpxB-dependent H_2_O_2_ production was the main antagonistic factor. Addition of *S. mitis* within hours after *S. mutans* inoculation was effective at reducing biofilm biomass, but not for 24 h pre-formed biofilms. Transcriptome analysis revealed responses for both *S. mitis* and *S. mutans*, with several *S. mutans* differentially expressed genes following a gene expression pattern previously described, while others being unique to the interaction with *S. mitis*. Finally, we show that *S. mitis* also affected coculture biofilm formation of several other commensal streptococci. Our study shows that strains with abundant H_2_O_2_ production are effective at inhibiting initial growth of caries pathogens like *S. mutans*, but are less effective at disrupting pre-formed biofilms and have the potential to influence the stability of other oral commensal strains.

## INTRODUCTION

Human dental caries (tooth decay) remains the most prevalent chronic disease in both children and adults, with 46% of children aged 2-19 years old experiencing active carious lesions as of 2019 (1). Decay is driven by the accumulation of acidic end products produced by oral bacteria within biofilms attached to the tooth surface as a byproduct from fermentation of dietary carbohydrates consumed by the host (2). *Streptococcus mutans*, and other mutans group streptococci species such as *Streptococcus sobrinus*, are most commonly associated with dental caries due to their displayed acidogenic and aciduric properties (3). Outgrowth of these cariogenic organisms can lead to an ecological shift within the microbial community, displacing commensals that are more sensitive to acidic environmental conditions (4). This perturbation then shifts the niche to favor outgrowth for additional acidogenic and aciduric species to persist and thrive, continuing the cycle and eventual development of disease (caries) (5, 6). Thus, thwarting the appearance of mutans group streptococci while keeping protective commensals intact remains a favorable therapeutic strategy towards caries prevention and overall maintenance of good oral health.

Oral commensals posse several mechanisms to combat the emergence of cariogenic bacteria. One is the arginine deiminase (ADS) pathway, which catalyzes the conversion of arginine to ornithine, ammonium, and carbon dioxide, while generating ATP from ADP and phosphate. The ammonia generated from this pathway is a critical factor for pH homeostasis, neutralizing the cytoplasm and alkalinizing the surroundings that offsets environmental acidification (7). A more direct mechanism to contest cariogenic bacteria is the production of hydrogen peroxide (H_2_O_2_). Exposure to H_2_O_2_ generates breakdown products of hydroxyl radicals and superoxide anions that can cause irreversible cellular damage in DNA integrity, oxidation of sulfurous amino acids and metal-binding sites within proteins, and mismetallation of enzymes (8). H_2_O_2_ is generated in part by a group of peroxigenic commensal streptococci via oxidase enzymes such as pyruvate oxidase (SpxB, Pox), which converts pyruvate to acetyl phosphate (9). Production of H_2_O_2_ by commensal streptococci such as *Streptococcus sanguinis* has been shown to inhibit the growth of *S. mutans* (10), and recently a strain of *Streptococcus oralis*, J22, was shown to disrupt *S. mutans* virulence both in an *ex vivo* tooth surface as well as reduced caries development within a rodent caries model (11). Therefore, abundant H_2_O_2_ production, which can be strain specific, has been a desirable characteristic of isolated oral bacteria that are screened and selected as potential probiotic candidates to prevent dental caries (12, 13). For example, the recent characterized isolate A12, recovered from the supragingival dental plaque of a caries-free individual, was able to both express the ADS pathway at high levels while also producing copious amounts of H_2_O_2_ that were sufficient to inhibit the growth of *S. mutans* (14, 15). Further identifying specific strains with these properties will not only expand our potential to develop probiotic candidates for use in a therapeutic setting, but additionally offer insights into how and why these strains are able to generate larger amounts of H_2_O_2_ compared to other strains of the same species through both molecular and genetic characterizations.

During a recent study on how human saliva modifies the behavior of oral streptococci (16), an eight species panel was tested for specific growth and biofilm formation phenotypes. During these experiments, we noted that one strain in particular, *Streptococcus mitis* ATCC 49456, more effectively inhibited the growth of *S. mutans* than other strains within the panel. The focus of this study was to further investigate how this growth inhibition originates, and whether it could be utilized to prevent both the emergence of, as well as disrupt pre-formed biofilms of, mutans group streptococci *S. mutans* and *S. sobrinus*. Our results reveal that this strain of *S. mitis* produces higher levels of H_2_O_2_ in a pyruvate oxidase-dependent manner, both in comparison to other strains of the *S. mitis* species, as well as other commonly studied strains of commensal streptococci. These results reinforce previous observations of the heterogeneity that exists between strains for H_2_O_2_ production, and its role in the inhibition of growth for cariogenic organisms of the oral cavity such as *S. mutans*.

## RESULTS

### Coculture with Streptococcus mitis inhibits biofilm formation of Streptococcus mutans

We cocultured *S. mutans* UA159 with seven different species of oral streptococci in 24 h biofilms and determined formed biomass using the crystal violet (CV) biofilm assay as a way to determine the level of antagonism between species **(Figure 1A)**. In all wells, some biomass in the cocultures could be visualized, except for wells that contained *S. mitis* ATCC 49456. CV remaining from the assay could not be quantified over background with *S. mitis*, but was detectable in all other cocultures tested, even at low levels (∼5-10% biomass remaining compared to *S. mutans* monoculture) as was the case for cocultures that included *S*. sp A12 and *Streptococcus gordonii* DL1 **(Figure 1B)**. We then visualized these cocultured biofilms by widefield microscopy. *S. mutans* formed characteristic rotund microcolony structures in the absence of *S. mitis* **(Figure 1C)**, but very little microcolony structure was observed in coculture **(Figure 1D)**.

**Figure 1.**
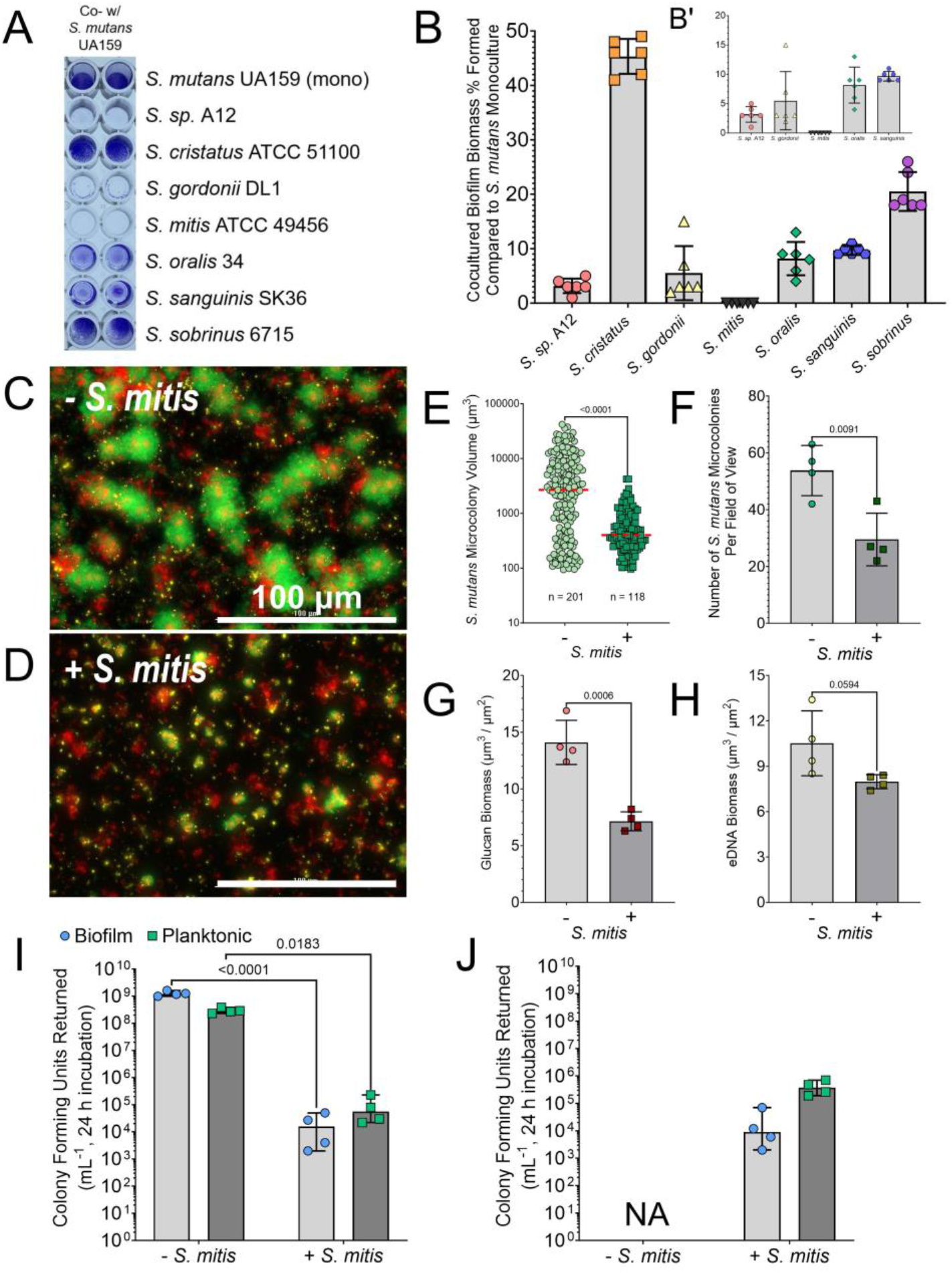
S. mitis Inhibits S. mutans Biofilm Formation. **(A)** Representative image of a crystal violet (CV) biofilm biomass assay where *S. mutans* is cocultured with different oral streptococci species (listed down right y-axis). *S. mutans* monoculture (mono) is shown at the top for reference. **(B)** CV quantification (Abs 575 nm) of the experiment shown in A. Data is expressed as the percentage of biomass formed in comparison to *S. mutans* monoculture (i.e., monoculture values set to 100%). **B’** is same data on a smaller y-axis with *S. cristatus* and *S. sobrinus* coculture data removed. n = 6. **(C)** Merged representative maximum intensity 40X Z-projection of 24 h *S. mutans* monoculture biofilm. *S. mutans* constitutively expresses GFP (green), eDNA was probed with labeled antibodies (yellow), and glucans visualized with labeled dextran (red). Scale bar (100 µm) is shown in the bottom right corner. **(D)** Merged representative maximum intensity 40X Z-projection of 24 h *S. mutans* cocultured biofilm with *S. mitis*. **(E)** Quantification of individual *S. mutans* microcolony volumes, **(F)** number of *S. mutans* microcolonies per field of view, **(G)** glucan biomass, and **(H)** eDNA biomass from the microscopy data shown in C and D. n = 4. Quantification was completed using Gen5 Image+ software. **(I)** *S. mutans* colony forming units (CFUs) returned from 24 h biofilms, with enumeration of cells in either biofilm or planktonic growth phase, grown with or without *S. mitis*. n = 4. **(J)** *S. mitis* CFUs returned. Data graphing and two-way analysis of variance (ANOVA) with multiple comparisons or student’s T-Test was completed in GraphPad Prism software.

The average *S. mutans* microcolony volume was only 495 μm^3^ in the presence of *S. mitis*, compared to 6013 μm^3^ in monoculture **(Figure 1E)**. In addition, there was a significant decrease in the number of *S. mutans* microcolonies formed per field of view (from 54 on average to 30) **(Figure 1F)**, and half of the glucan biomass was present (14 to 7 μm^3^/μm^2^) **(Figure 1G)**, while eDNA biomass was also generally decreased with addition of *S. mitis*, but not statistically significant **(Figure 1H)**. These data show that coculture with *S. mitis* dramatically reduces the amount of *S. mutans* biofilm formed in a 24 h period.

We next enumerated the colony forming units (CFUs) returned for *S. mutans* with and without addition of *S. mitis* to determine if the biofilm phenotype displayed was due to general growth inhibition or from *S. mutans* cells failing to enter into a biofilm state. Cells recovered from biofilm supernatants (i.e., planktonic cells) were plated in comparison to adhered cells within a biofilm. Only 1×10^5^ CFU mL^-1^ of *S. mutans* cells were returned from either biofilm or planktonic populations with addition of *S. mitis*, which was equivalent to our starting inoculum **(Figure 1I)**. This suggests that the biofilm phenotype is from general growth inhibition due to the presence of *S. mitis*. In addition, *S. mitis* also sparingly grew in the coculture with only a log increase in CFUs returned over the course of the experiment **(Figure 1J)**.

### S. mutans growth inhibition is specific for S. mitis ATCC 49456

To ensure that our loss of biofilm biomass was applicable to all strains of *S. mutans* and not specific for UA159 (SMU159), we cocultured 19 other isolates of *S. mutans* with *S. mitis* **(Figure 2A)**. All isolates tested showed a similar abolishment of biofilm formation with *S. mitis*. Similarly, we also wanted to determine if all strains of *S. mitis* could inhibit *S. mutans* growth. Only strain ATCC 49456, out of four total *S. mitis* strains tested, completely eliminated *S. mutans* biofilm biomass accumulation **(Figure 2B)**. Coculture with strains B6, SK306 and SK569 amassed significantly more quantifiable biomass than with ATCC 49456, and were at levels even higher than that of strain *S. oralis* 34 **(Figure 2C)**. Microscopy with these three strains revealed observable microcolony formation of *S. mutans* **(Figure 2D)**, which translated to significantly higher quantifiable *S. mutans* microcolony volumes, biofilm thickness (i.e., average microcolony height), and glucan biomass **(Figures 2E, 2F and 2G)**. In all, these data show that *S. mutans* growth inhibition is specific to ATCC 49456 out of the *S. mitis* strains tested here and is a general phenotype to all *S. mutans* isolates assayed.

**Figure 2.**
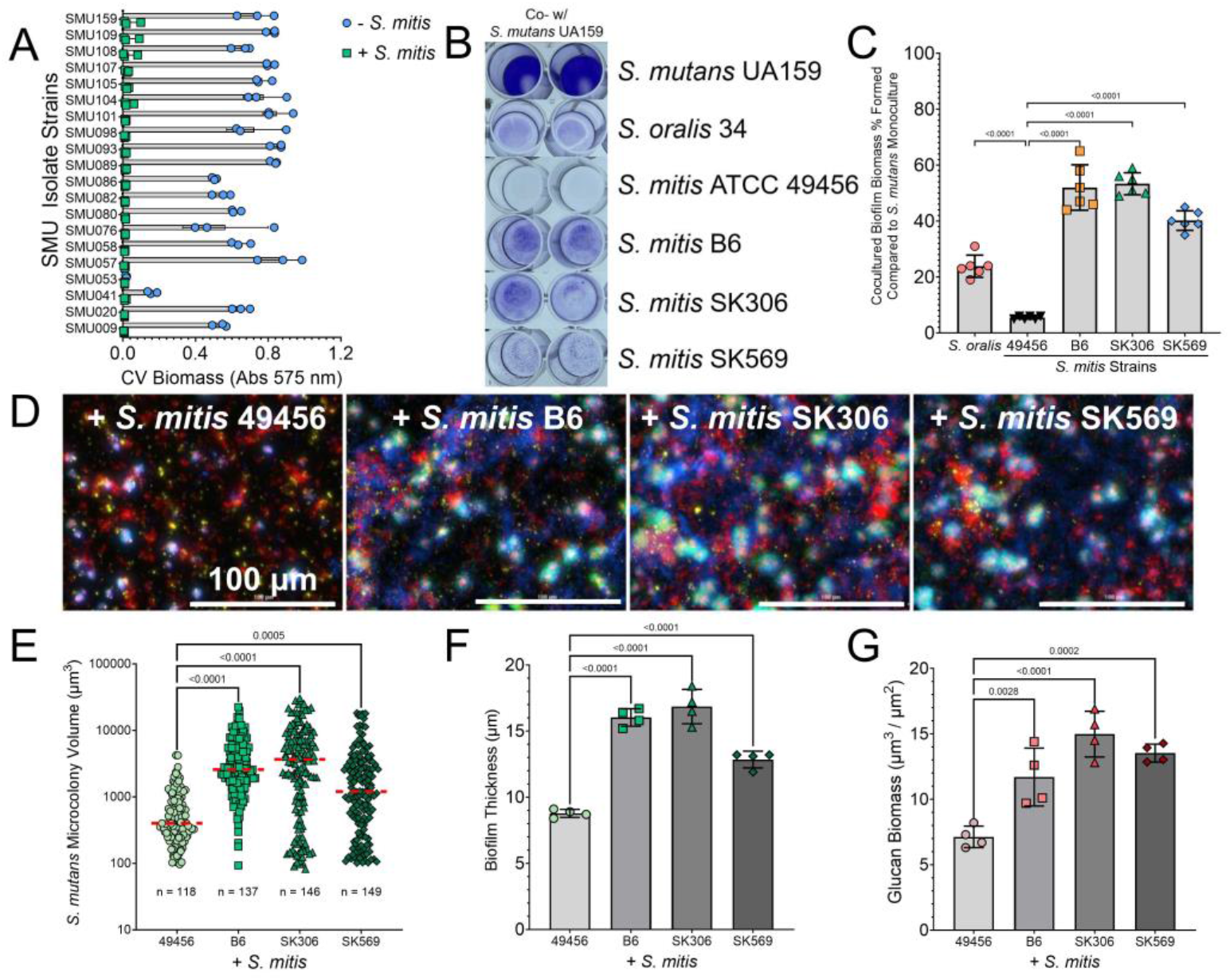
Strain ATCC 49456 Specifically Inhibits S. mutans Biofilm Formation. **(A)** CV quantification (Abs 575 nm) of biomass formed by various *S. mutans* isolates grown in monoculture (-*S. mitis*) or in coculture with *S. mitis* (+ *S. mitis*). **(B)** Representative image of a CV biofilm biomass assay where *S. mutans* is cocultured with different strains of *S. mitis* (listed down right y-axis). Coculture with *S. oralis* is included as a reference. **(C)** CV quantification (Abs 575 nm) of the experiment shown in B. Data is expressed as the percentage of biomass formed in comparison to *S. mutans* monoculture. **(D)** Merged representative maximum intensity 40X Z-projection of 24 h *S. mutans* cocultured biofilms with different strains of *S. mitis. S. mutans* constitutively expresses GFP (green), eDNA was probed with labeled antibodies (yellow), glucans visualized with labeled dextran (red), and a total cell strain applied to visualize *S. mitis* within the biofilms (Hoechst 33342; blue). Scale bar (100 µm) is shown in the bottom right corner. **(E)** Quantification of individual *S. mutans* microcolony volumes, **(F)** biofilm thickness, and **(G)** glucan biomass from the microscopy data shown in D. n = 4. Quantification was completed using Gen5 Image+ software. Data graphing and one-way analysis of variance (ANOVA) with multiple comparisons was completed in GraphPad Prism software.

### S. mitis ATCC 49456 produces high levels of hydrogen peroxide

To identify factor(s) that may be involved in growth inhibition of *S. mutans*, we first tested whether spent supernatants of *S. mitis* ATCC 49456 could inhibit growth of *S. mutans* to a similar degree as direct inoculation. Cultures of *S. mitis* alone or *S. mitis* cocultured with *S. mutans* were grown for 24 h prior to supernatants being removed after centrifugation, along with supernatants of *S. mutans* and *S. gordonii* grown in monoculture as controls **(Figure 3A)**. While all tested supernatants displayed a decrease in *S. mutans* biomass compared to fresh medium, none including those with *S. mitis*, was similar to direct *S. mitis* inoculation **(Figure 3B)**. Thus, we ruled out specific production of a factor(s), such as bacteriocin(s), that could accumulate in the supernatant of *S. mitis* to inhibit *S. mutans* growth.

**Figure 3.**
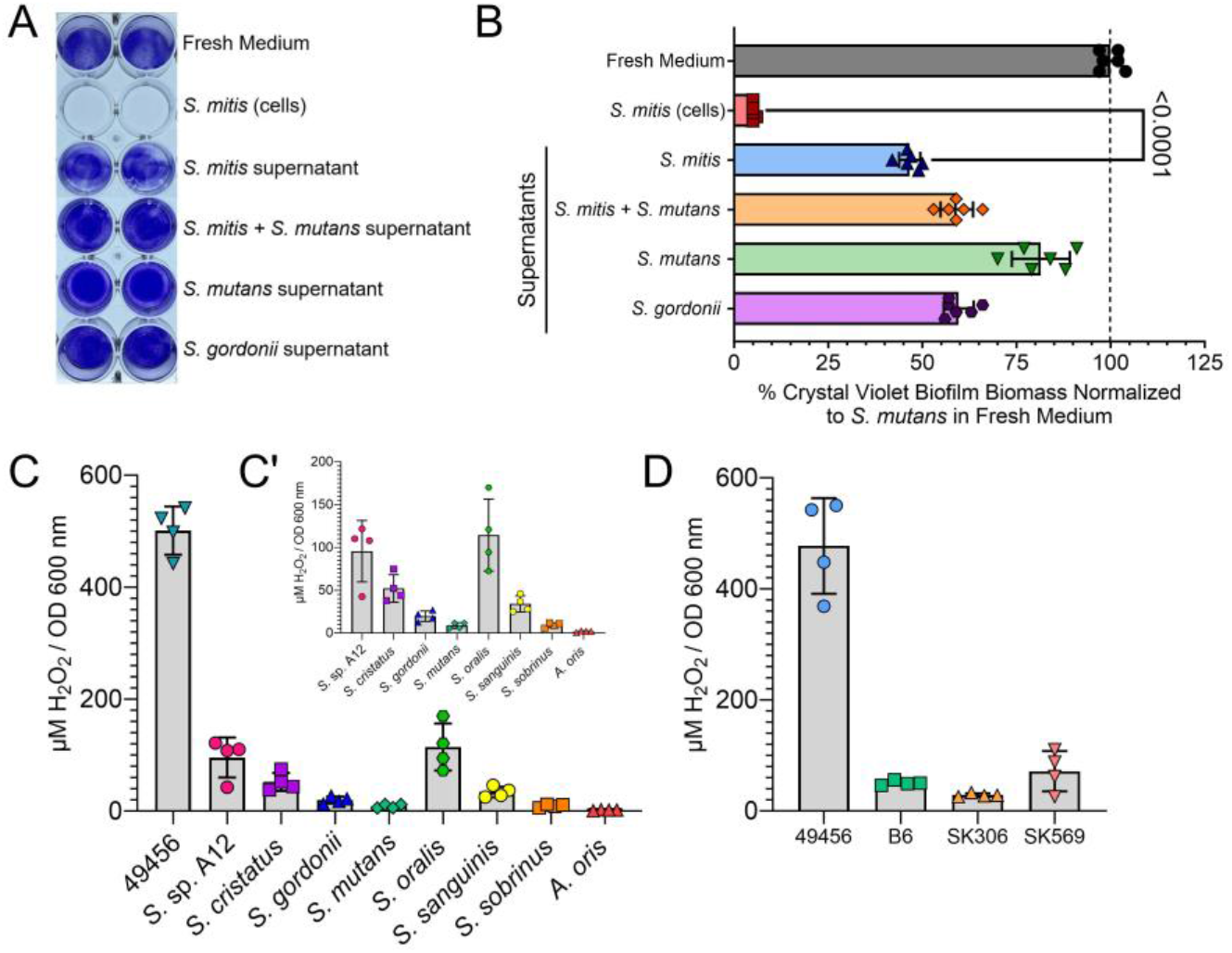
S. mitis ATCC 49456 Produces High Levels of Hydrogen Peroxide. (A) Representative image of a crystal violet (CV) biofilm biomass assay where *S. mutans* is grown in fresh medium, cocultured with *S. mitis* (cells), or grown in extracted 24 h supernatants of various cultures (listed down right y-axis). **(B)** CV quantification (Abs 575 nm) of the experiment shown in A. Data is expressed as the percentage of biomass formed in comparison to *S. mutans* grown in fresh medium (i.e., monoculture values set to 100%). n = 6. **(C)** Quantification of hydrogen peroxide present in culture supernatants of different oral species. Values were normalized to culture density (OD_600 nm_) prior to centrifugation and extraction of culture supernatants. *S. mitis* (ATCC 49456) is shown on the left. **C’** is same data on a smaller y-axis with *S. mitis* data removed. n = 4. **(D)** Quantification of hydrogen peroxide present in culture supernatants of different *S. mitis* strains. Data graphing and one-way analysis of variance (ANOVA) with multiple comparisons was completed in GraphPad Prism software.

A common way for other oral streptococci to antagonize *S. mutans* is through hydrogen peroxide (H_2_O_2_) production (17). We collected overnight supernatants from our original oral streptococci panel (Figure 1A) along with *Actinomyces oris* as a control (non-H_2_O_2_ producing strain) and quantified the amount of H_2_O_2_ present normalized to culture optical density at the time of harvest **(Figure 3C)**. We found *S. mitis* ATCC 49456 to produce 4-5 fold higher amounts of H_2_O_2_ compared to our next highest species (*S*. sp. A12 and *S. oralis* 34), which was notable on Prussian blue agar plates as well **(Supplemental Figure 1)**. In addition, ATCC 49456 produced 5-18 fold higher amounts of H_2_O_2_ compared to *S. mitis* strains B6, SK306, and SK569 **(Figure 3D)**. From these findings, we reasoned that this high amount of H_2_O_2_ production was the unknown factor(s) leading to growth inhibition of *S. mutans* in biofilm populations.

### S. mitis ATCC 49456 inhibits S. mutans through hydrogen peroxide production

To confirm the role of *S. mitis* ATCC 49456 H_2_O_2_ production in the growth inhibition of *S. mutans*, we first confirmed that addition of 100 U mL^-1^ catalase to the growth medium, as well as mutation of the pyruvate oxidase gene (*spxB*) in *S. mitis* ATCC 49456, almost eliminated the amount of detectable H_2_O_2_ production in the recovered culture supernatant **(Figure 4A)**. We then tested these conditions in a coculture CV biofilm assay between *S. mutans* and *S. mitis* ATCC 49456 **(Figure 4B)**. Notably, addition of catalase resulted in 60% biofilm biomass remaining compared to *S. mutans* monoculture and 34% for the coculture containing the *spxB* mutant – a significant increase from the 1% observed with the *S. mitis* wild-type (WT) strain lacking catalase **(Figure 4C)**. Substantial differences were also observed in imaging of the biofilms, as more *S. mutans* biomass could be visualized in conditions containing catalase or Δ*spxB* **(Figure 4D)**. This was confirmed by quantifying the images and measuring significant increases in *S. mutans* microcolony volumes **(Figure 4E)**, *S. mutans* biofilm thickness **(Figure 4F)**, and increases in detectable glucan biomass **(Figure 4G)**. It is important to note that catalase addition did not affect *S. mutans* biofilm formation in monoculture **(Supplemental Figure 2)**. In all, these data confirm that the robust H_2_O_2_ production by *S. mitis* ATCC 49456 was largely responsible for the biofilm phenotype observed in coculture with *S. mutans*.

**Figure 4.**
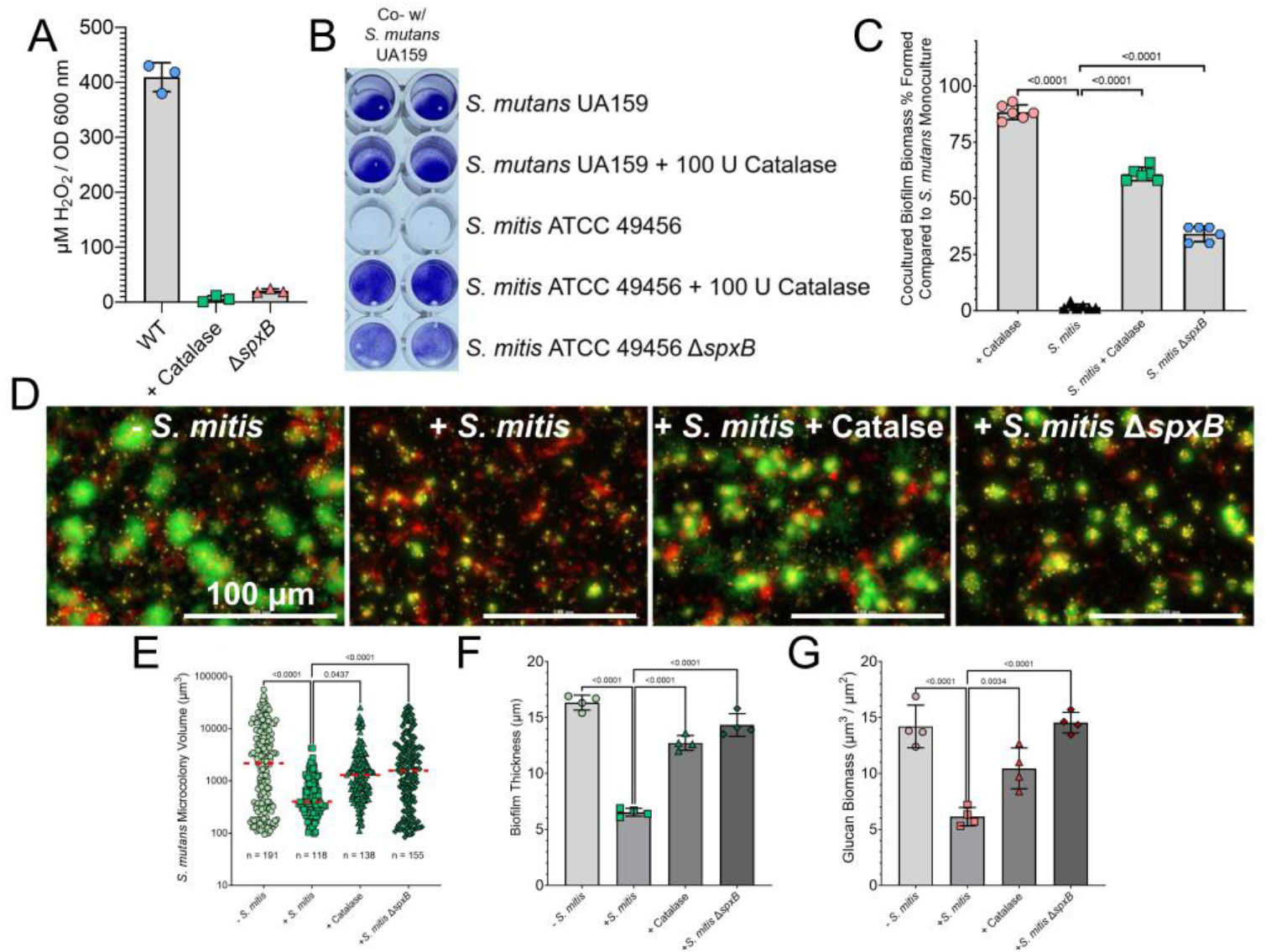
S. mitis ATCC 49456 Hydrogen Peroxide Production Inhibits S. mutans Biofilm Formation. **(A)** Quantification of hydrogen peroxide present in culture supernatants of *S. mitis* ATCC 49456 wild-type (WT), with addition of 100 U mL^-1^ catalase to the growth medium (+ Catalase), and with an *spxB* mutant (Δ*spxB*). n = 3. **(B)** Representative image of a CV biofilm biomass assay where *S. mutans* is cultured in different conditions (listed down right y-axis). **(C)** CV quantification (Abs 575 nm) of the experiment shown in B. Data is expressed as the percentage of biomass formed in comparison to *S. mutans* monoculture alone (i.e., monoculture values set to 100%). n = 6. **(D)** Merged representative maximum intensity 40X Z-projection of 24 h *S. mutans* biofilms grown in monoculture (-*S. mitis*), in coculture with *S. mitis* (+ *S. mitis*), with *S. mitis* and addition of 100 U mL^-1^ catalase (+ *S. mitis* + Catalase), and with the *S. mitis spxB* mutant (+ *S. mitis* Δ*spxB*). *S. mutans* constitutively expresses GFP (green), eDNA was probed with labeled antibodies (yellow), and glucans visualized with labeled dextran (red). Scale bar (100 µm) is shown in the bottom right corner. **(E)** Quantification of individual *S. mutans* microcolony volumes, **(F)** biofilm thickness, and **(G)** glucan biomass from the microscopy data shown in D. n = 4. Quantification was completed using Gen5 Image+ software. Data graphing and one-way analysis of variance (ANOVA) with multiple comparisons was completed in GraphPad Prism software.

### S. mitis ATCC 49456 disrupts forming but not pre-formed biofilms of caries pathogens

To this point, addition of *S. mitis* always occurred during inoculation of *S. mutans* biofilms (i.e., t = 0 h). To determine if *S. mitis* ATCC 49456 could disrupt biofilms of caries pathogens at different stages of development, we performed a CV biofilm assay where the growth medium was removed from incubating *S. mutans* or *S. sobrinus* monoculture biofilms at different hours post inoculation and replaced with either fresh medium (FM) or medium that contained *S. mitis* **(Figure 5A)**. Less than 10% of biomass in the +*S. mitis* condition compared to FM only was documented when *S. mitis* was added during the first four hours of incubation for *S. mutans*, but more biofilm biomass was present at time points 5, 6 and 7 hours **(Figure 5B)**. Only a 14% reduction in biomass with *S. mitis* was recorded when *S. mitis* as added after 7 hours. A change in accumulated biomass with prolonged *S. mitis* addition was less drastic with *S. sobrinus* 6715, but followed a similar trend. Next, we attempted to disrupt *S. mutans* biofilms that had been incubated for 24 h prior to *S. mitis* ATCC 49456 addition. We replaced the medium of biofilms with either 1X PBS (corresponding to biomass after 24 h), FM or medium containing low, medium or high optical densities of *S. mitis* WT or Δ*spxB* **(Figure 5C)**. There were no significant differences observed between WT and Δ*spxB* at the high optical density inoculation, but differences were present with the medium and low optical densities **(Figure 5D)**. However, 82% of the biomass remained with the medium density, compared to the 1X PBS control, and 103% with the low density *S. mitis* WT. We also observed *S. mutans* microcolony architecture 24 h post *S. mitis* addition **(Figure 5E)**. Biofilms that were incubated with the Δ*spxB* strain actually displayed less biomass than the WT strain at all added densities **(Figure 5F)**. Therefore, these data suggest that while *S. mitis* ATCC 49456 can effectively inhibit forming biofilms, coculture is not effective at disrupting pre-formed biofilms in a H_2_O_2_-dependent manner.

**Figure 5.**
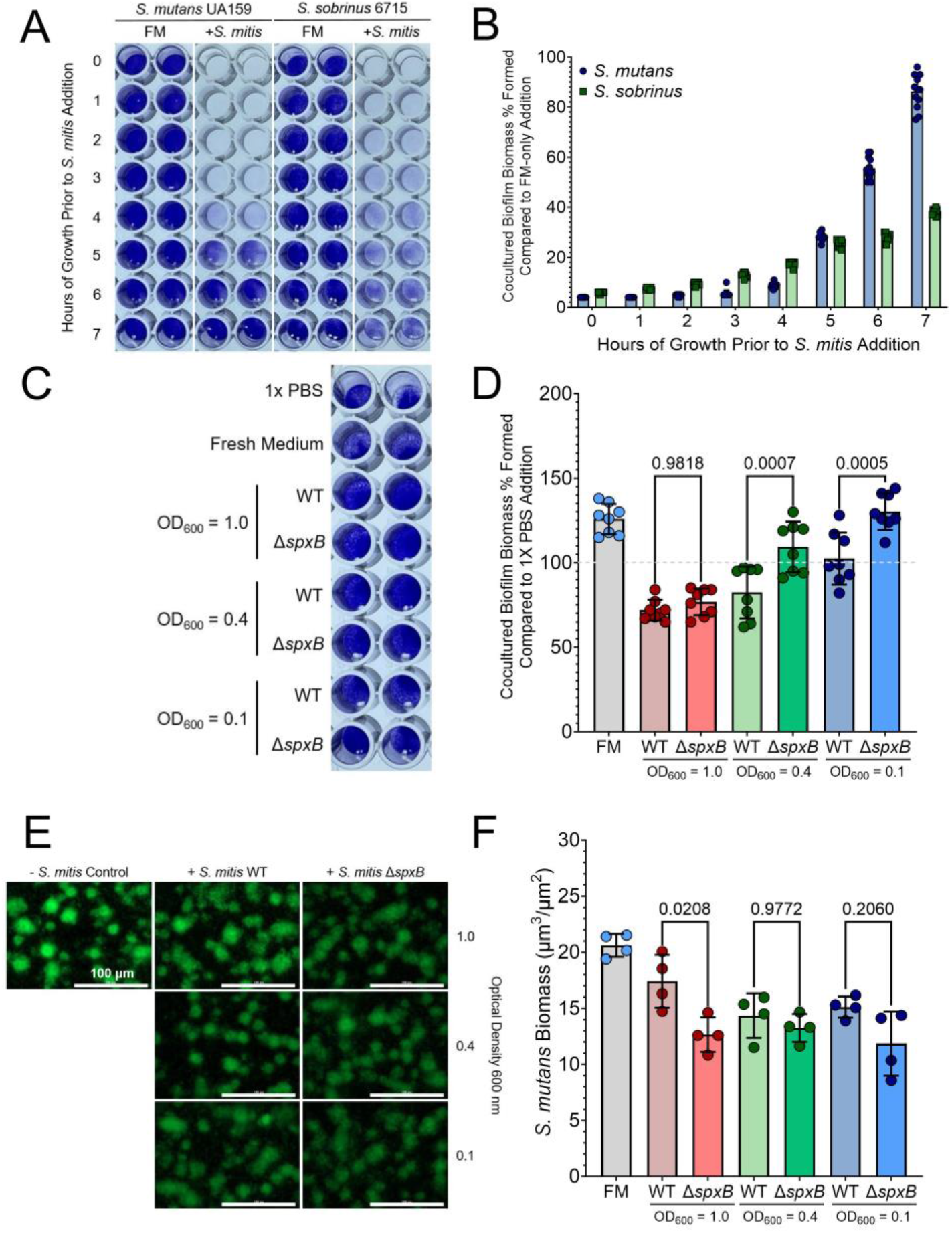
S. mitis Disruption of Forming and Pre-formed Biofilms of Caries Pathogens. **(A)** Representative image of a CV biofilm biomass assay where either fresh medium (FM) or medium containing *S. mitis* ATCC 49456 replaced the original growth medium of either *S. mutans* or *S. sobrinus* biofilms at the time point indicated on the left y-axis. 0 indicates addition during the inoculation of *S. mutans* or *S. sobrinus*. Biofilms were grown for a total of 24 h. **(B)** CV quantification (Abs 575 nm) of the experiment shown in A. Data is expressed as the percentage of biomass remaining in the *S. mitis* coculture condition compared to FM addition at each specific time point. **(C)** Representative image of a CV biofilm biomass assay where medium from 24 h pre-formed *S. mutans* biofilms is replaced with either 1X PBS, FM, or *S. mitis* ATCC 49456 wild-type (WT) or *spxB* mutant (Δ*spxB*) at different optical densities (OD_600 nm_ = 1.0, 0.4, or 0.1). The biofilms were then grown for another 24 h prior to CV staining. **(D)** CV quantification (Abs 575 nm) of the experiment shown in C. Data is expressed as the percentage of biomass remaining in comparison to the 1X PBS control (i.e., biofilm formed at 24 h without additional growth). FM refers to the addition of fresh medium (i.e., no *S. mitis* added). n = 8. **(E)** Representative maximum intensity 40X Z-projection of 48 h *S. mutans* biofilms grown in the absence of (-*S. mitis*), or with the addition of, *S. mitis* WT or Δ*spxB* at different optical densities at 24 h. Biofilms were then grown for another 24 h prior to imaging. *S. mutans* constitutively expresses GFP (green). Scale bar (100 µm) is shown in the bottom right corner. **(F)** Quantification of *S. mutans* biomass from the microscopy data shown in E. FM refers to the – *S. mitis* control (i.e., no *S. mitis* added). n = 4. Quantification was completed using Gen5 Image+ software. Data graphing and two-way analysis of variance (ANOVA) with multiple comparisons was completed in GraphPad Prism software.

### Transcriptome profiling of S. mitis and S. mutans cocultures

Previously our group has explored how *S. mutans* alters its behavior in the presence of various oral bacteria in varying environmental conditions through RNA-Seq (16, 18). To determine how *S. mutans* responds to a high H_2_O_2_ producer such as *S. mitis* ATCC 49456, we harvested mono- and cocultures of both species at mid-exponential log phase and extracted RNA to profile transcriptomes when growing alone versus in competition against each other. We documented only 31 differentially expressed genes (DEGs) in *S. mitis* coculture compared to monoculture, with 29 upregulated and only 2 downregulated **(Figure 6A and 6B, Supplemental Table 1)**. The 29 upregulated DEGs consisted of genes involved in carbohydrate utilization, including an operon consisting of ABC transporter permeases (SM12261_RS02685 - SM12261_RS02705), alcohol dehydrogenases, and dihydroxyacetone kinase subunits L and M (*dhaL* and *dhaM*) **(Figure 6C)**. Interestingly, the two downregulated DEGs for *S. mitis* were an annotated bacteriocin immunity protein (SM12261_RS06165) and the histidine kinase *comD* **(Figure 6D)**. In contrast, a total of 212 DEGs were found for *S. mutans* in coculture with *S. mitis* **(Figure 6E and 6F, Supplemental Table 2)**. Of the 134 upregulated DEGs, several corresponded to mobile genetic elements and the genomic islands TnSmu1 and TnSmu2, ABC transporters for iron (SMU_995 – SMU_998) and amino acids (SMU_932 – SMU_936), ribosomal proteins and unknown hypotheticals **(Figure 6G)**. 78 downregulated DEGs consisted of genes related to pyrimidine biosynthesis, bacteriocins, as well as PTS systems for fructose (SMU_872), fructose/mannose (SMU_1956c – SMU_1961c), trehalose (SMU_2037 and SMU_2038) and *manLMN* **(Figure 6H)**.

**Figure 6.**
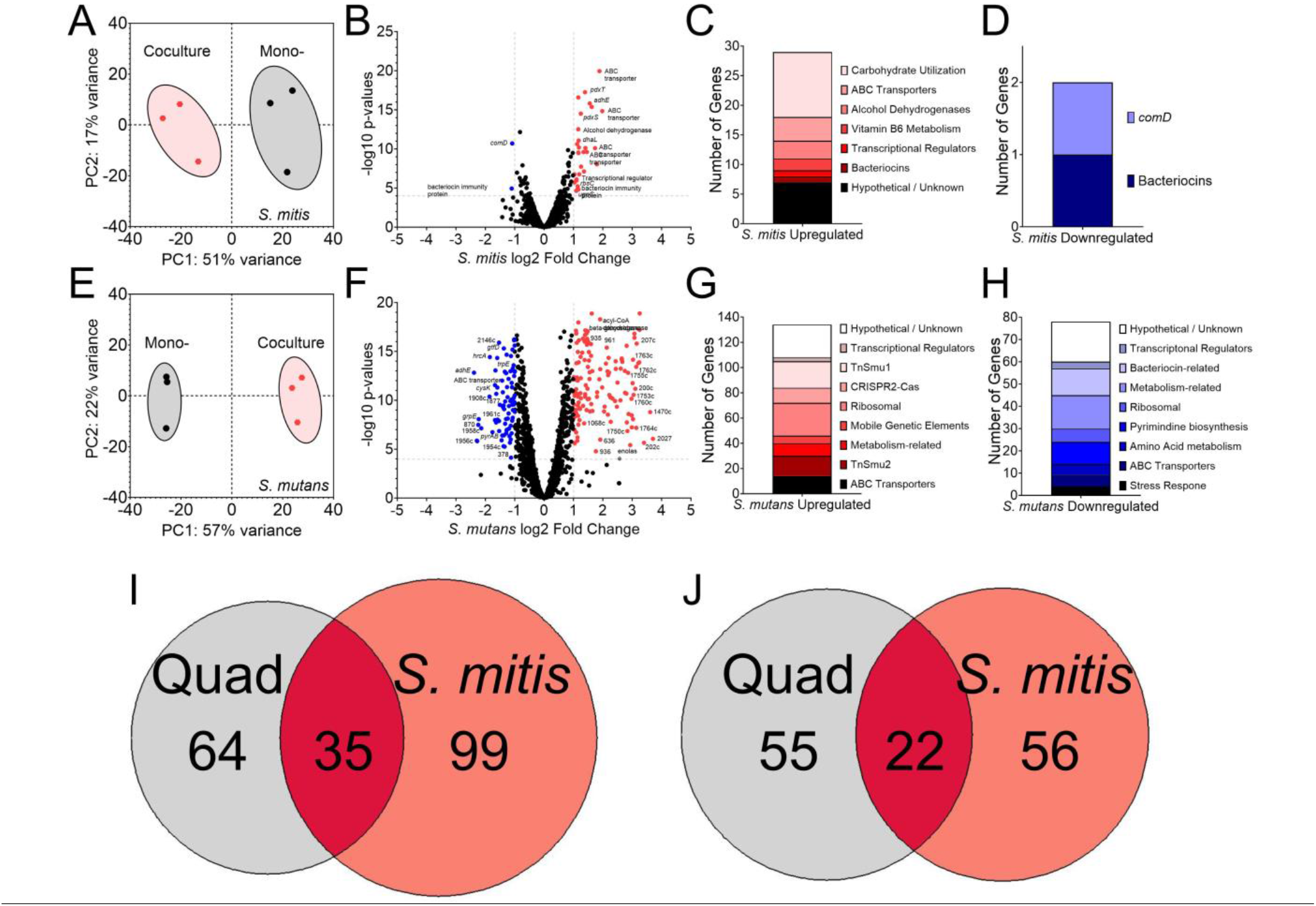
Transcriptomes of S. mutans and S. mitis ATCC 49456 during Coculture Growth. **(A)** Principal component analysis (PCA) from RNA-Seq expression data (n = 3) of *S. mitis* grown in monoculture (black circles) or coculture with *S. mutans* (red hexagons). The proportion of variance for either PC1 (x-axis) or PC2 (y-axis) are listed. **(B)** Volcano plot of changes within individual *S. mitis* genes (circles) between monoculture and coculture with *S. mutans*. Differentially expressed genes (DEGs; = genes with ≥4 Log10 *p* value and Log2 fold change ≥ (-)1) are shown in either red (upregulated, right) or blue (downregulated, left). Individual gene identifier, name and/or characterized function are displayed, if able. **(C)** Stacked bar chart of upregulated *S. mitis* DEGs from the dataset grouped by pathway/operon/function. **(D)** Stacked bar chart of downregulated *S. mitis* DEGs. **(E)** PCA from RNA-Seq expression data of *S. mutans* grown in monoculture (black circles) or coculture with *S. mitis* (red hexagons) (n = 3). **(F)** Volcano plot of changes within individual *S. mutans* genes between monoculture and coculture with *S. mitis*. **(G)** Stacked bar chart of upregulated *S. mutans* DEGs from the dataset grouped by pathway/operon/function. **(H)** Stacked bar chart of downregulated *S. mutans* DEGs. **(I)** Venn diagram of the number of upregulated *S. mutans* DEGs from growth in quadculture (included *S. gordonii, S. oralis* and *S. sanguinis*; grey) or coculture with *S. mitis* (red). **(J)** Venn diagram of the number of downregulated *S. mutans* DEGs. Data graphing and PCA calculations were completed in GraphPad Prism software.

To gain a better understanding of how *S. mutans* may specifically respond to *S. mitis* ATCC 49456, we compared our DEGs found here to a previous list of DEGs we documented when *S. mutans* was in quadculture with *S. gordonii, S. oralis* and *S. sanguinis* (18). Of the 198 total upregulated DEGs found between both studies, only 35 were shared between the two datasets and 99 upregulated DEGs were specific to the interaction with *S. mitis* **(Figure 6I and Supplemental Table 3)**. Common upregulated genes included the CRISPR2-Cas system (SMU_1750c – SMU_1764c), genes within TnSmu1, and the transcriptional regulator *hdiR* (SMU_2027). Several of the *S. mitis*-specific upregulated genes were within operons of the common upregulated genes (e.g., genes within TnSmu1 and ribosomal proteins), while others were clearly specific to the interaction with *S. mitis* such as the ABC transporters for iron and amino acids, as well as genes within TnSmu2. For downregulated DEGs, genes specific for the quadculture vs *S. mitis* were almost evenly split 55 to 56 with 22 shared between both out of the total of 133 genes **(Figure 6J and Supplemental Table 4)**. Here, downregulated genes shared mainly related to *S. mutans* bacteriocins, the fructose/mannose PTS (SMU_1956c – SMU_1961c), and *cysK*. Downregulation of *manLMN* and the trehalose PTS was specific to *S. mitis*, while the TnSmu2 genomic island was downregulated in our previous quadculture compared to upregulated here with *S. mitis*.

### Growth with S. mitis impairs biofilm formation of other oral streptococci

Aside from growth with caries pathogens *S. mutans* and *S. sobrinus*, we wanted to determine if *S. mitis* ATCC 49456 also affected the biofilm formation of other oral streptococci **(Figure 7A)**. Indeed, coculturing with *S. mitis* reduced the biomass of *S. cristatus* ATCC 51100 by over 50%, and close to 50% with *S. sanguinis* SK36 as well **(Figure 7B)**. Coculturing with other streptococci, such as *S. gordonii* DL1 or *S. oralis* 34 only had marginal reductions of biomass of around 20%. While the biofilm inhibition phenotype displayed by *S. mitis* ATCC 49456 is most pronounced with caries pathogens *S. mutans* and *S. sobrinus, S. mitis* ATCC 49456 can cause similar biofilm phenotypes with other oral streptococci to varying degrees.

**Figure 7.**
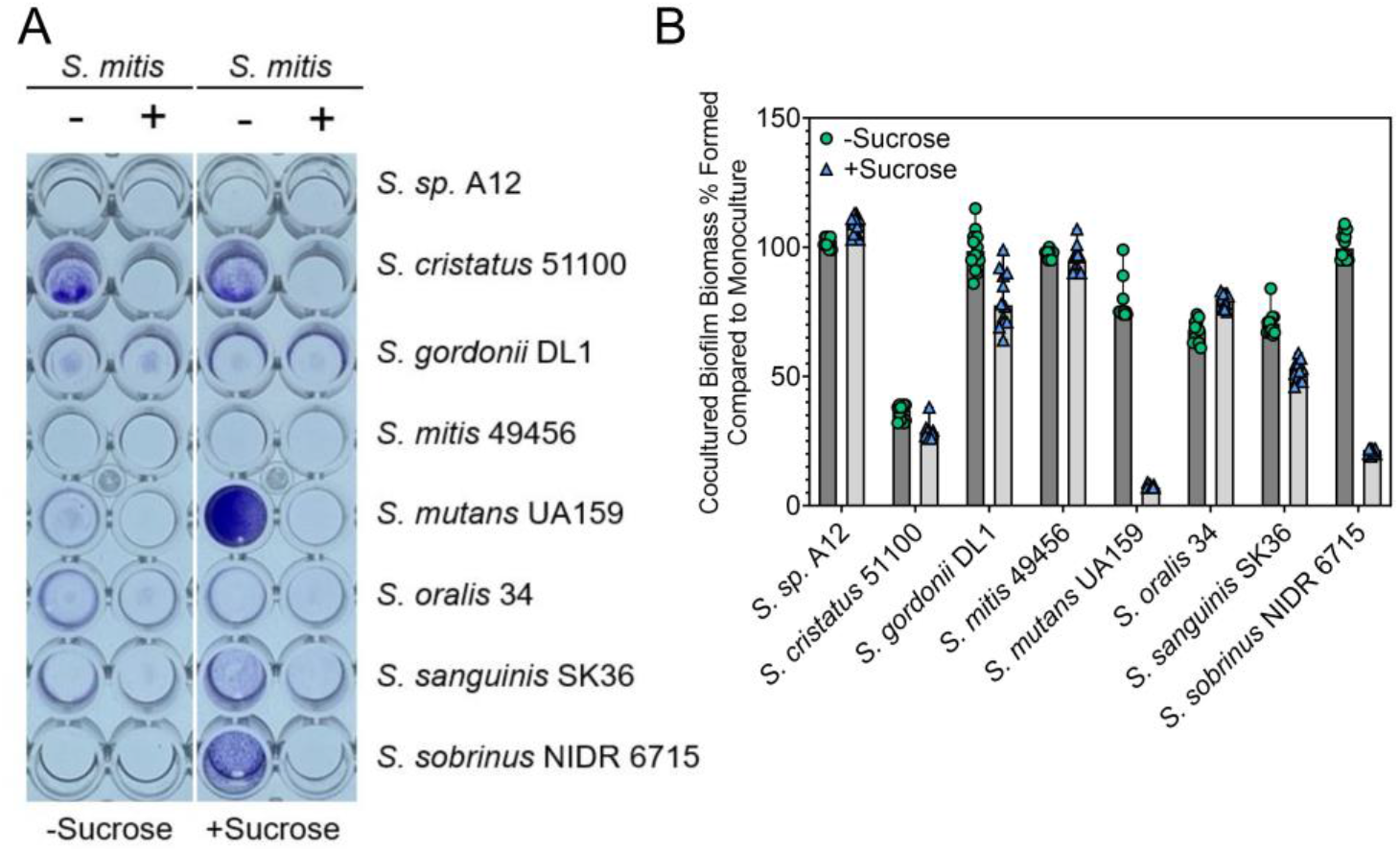
S. mitis ATCC 49456 Impacts Biofilm Formation of other Oral Streptococci. **(A)** Representative image of a CV biofilm biomass assay of different oral *Streptococcus* species listed on the right y-axis, grown in monoculture (-) or in coculture with *S. mitis* (+) in medium lacking (-Sucrose) or containing (+Sucrose) sucrose. **(B)** CV quantification (Abs 575 nm) of the experiment shown in A. Data is expressed as the percentage of biomass remaining in the *S. mitis* coculture condition compared to the monoculture condition, which lacks *S. mitis*. Dark grey bars with green circles represents TYG medium lacking sucrose (-Sucrose), while light grey bars with blue triangles represents TYG medium containing sucrose (+Sucrose). n = 12. Data graphing were completed in GraphPad Prism software.

## DISCUSSION

The oral streptococci that colonize the tooth and other surfaces in the oral cavity must contend for nutritional resources and physical space with other microbes within their niche to persist and thrive. Some oral streptococci form synergistic relationships with other species, as is the case with *S. mitis* ATCC 49456 and *Corynebacterium matruchotii* in corncob arrangements that are commonly seen in supragingival dental plaque (19, 20). At other times, these relationships are antagonistic, as observed in the interactions of oral commensal streptococci with *S. mutans*. It is well established that H_2_O_2_ production by peroxigenic oral streptococci is a major antagonistic mechanism that several cariogenic organisms such as *S. mutans* or *Actinomyces* are sensitive to (10, 17, 21), and a desirable trait for potential oral probiotic species (22). Our work here reinforces several previous findings that show strains that are high H_2_O_2_ producers can effectively outcompete *S. mutans* through growth inhibition (11, 14, 15). In addition, there is heterogeneity amongst strains of a given species in the amount of H_2_O_2_ produced – we recorded 5-18x higher H_2_O_2_ accumulated with ATCC 49456 compared to three other *S. mitis* strains that were available to us, including B6. Previously, Velsko *et al*. (13) showed in a panel of 113 oral streptococci clinical isolates that a high degree of variation presented in mean antagonism displayed against *S. mutans* in a spot-based agar plate competition assay, including within isolates from the same species. One possible source in the altered levels of antagonism seen is the ability of select isolates to produce higher amounts of H_2_O_2_, although a relationship between pyruvate oxidase gene phylogeny and antagonistic activity could not be determined in that study (13). Still other factors, such as regulation of *spxB* transcriptional activity between strains, could account for differences in H_2_O_2_ output. For example, *spxB* is influenced by carbohydrate availability in *S. gordonii* and is regulated in a CcpA-dependent manner, with expression being repressed in the presence of glucose (23, 24). All of our experiments here were conducted in medias containing glucose, opening the potential that some isolates break away from these common regulation paradigms, either through mutation(s) in regulators or in regulatory elements (i.e., transcriptional binding sites) that direct the promoter activity of *spxB*. Identifying specific strains of the same species that contrast in H_2_O_2_ production will assist in this exploration, as we have done here. Of note, several mutations are present in the *spxB* promoter region between strains B6 and ATCC 49456 that could be located within *cre* sites for CcpA binding **(Supplemental Figure 3)**, although the exact *cre* site sequences are difficult to discern without further experimentation. Another possibility is post-translational modification of SpxB, although the predicted sites of modification are conserved between the strains (data not shown) (25).

The role of H_2_O_2_ in antagonism of *S. mutans* is commonly tested with either both strains being inoculated together (as we initially have done in our coculture biofilm assays) or the H_2_O_2_ producer grown first prior to the inoculation/spotting of *S. mutans* (12). We took the opportunity with the phenotypes displayed by ATCC 49456 to test whether a highly antagonistic strain could disrupt pre-formed biofilms of *S. mutans*, a characteristic that would be desirable for a probiotic strain designed to be given to individuals at the initial stages of caries development (i.e., where *S. mutans* would be expected to be already present). We found the impact of added ATCC 49456 to 24 h biofilms of *S. mutans* to be minimal, even at different cell densities. The largest change in biomass, observed with addition of *S. mitis* at a high cell density, was not dependent on the presence of an intact *spxB* gene, suggesting the effect observed was not due to H_2_O_2_ production. One limitation of our experiments shown here is that we only quantified cell biomass through CV staining or biomass quantification of microscopy images – changes may have resulted in the number of viable cells remaining after *S. mitis* addition, which would be a desirable outcome. We attempted to address this with a live/dead staining of treated and control *S. mutans* biofilms (data not shown). However, we found that 70-80% of *S. mutans* UA159 cells stained “dead” in our untreated controls, making interpretations on the impact of *S. mitis* addition to these biofilms difficult, and thus was not included here. Still, our data suggests that an antagonistic strain such as ATCC 49456 would not be optimal to disturb pre-formed biofilms. The best strategy for addition of antagonistic strains remains prior to or during biofilm formation of caries pathogens. Our time course assay within 7 h of initial biofilm formation shows there is potential for an “intervention window” during the earliest stages of growth where an antagonistic strain could be added after *S. mutans*, but the window closes rapidly before the largest effects are lost (< 4 h after *S. mutans* inoculation).

Another area we briefly examined was the impact of ATCC 49456 on the biofilm formation of other oral streptococci. In addition to *S. mutans* and *S. sobrinus, S. mitis* also influenced the biofilm formation of commensals *S. cristatus* and *S. sanguinis*, with *S. gordonii* and *S. oralis* seeing moderate reductions as well. The impact of high H_2_O_2_ producers on other commensals and stability of the health-associated microbiome has not been fully investigated, but is relevant towards the usage of such strains as probiotic candidates. An ideal strain(s) would not significantly affect the persistence of other commensals within the shared niche. There is potential that the highest H_2_O_2_ producer that displays phenotypes similar to ATCC 49456 is not an optimal candidate. Rather, selected strain(s) that are higher than species average (e.g., strains in the 65^th^ – 85^th^ percentile), yet do not interfere with the stability of other niche members, could be more optimal choices. In addition, above average H_2_O_2_ accumulation could be paired with other qualities such as high ADS activity as seen with A12 (14), or production of bacteriocin-like peptides in the case of *S. oralis* subspecies *dentisani* (26), as a well-rounded probiotic candidate that could antagonize the growth of cariogenic organisms via multiple mechanisms. It is also important to note that ATCC 49456 failed to effectively adhere and form biofilms on its own in monoculture. In fact, ATCC 49456 (NCTC 12261) has been shown to have a strong site-tropism for the buccal mucosa over supragingival plaque where it would encounter caries pathogens such as *S. mutans* (27). One issue that has been encountered with potential probiotic candidates is their transient colonization of animal models where there competition against caries organisms like *S. mutans* is directly tested *in vivo* (28). Another quality to consider is the ability of these strains to form biofilms and attach to surfaces without the assistance of mutans group-derived glucans. With recent knowledge gained regarding the oral microbiome’s biogeography and that certain species of oral streptococci are site-specialists within a given niche (27, 29, 30), the search could be narrowed to focus on species such as *S. cristatus, S. gordonii, S. oralis* and *S. sanguinis* who are considered to be supragingival plaque specialists.

Finally, we compared the transcriptomes of ATCC 49456 and *S. mutans* growing in coculture versus alone in monoculture. Our group has previously documented a conserved gene expression pattern in *S. mutans* when cultured with other oral commensal streptococci but not with other disease-related oral streptococci (i.e., *S. sobrinus*) or non-*Streptococcus* oral bacteria in various conditions (16, 18). By expanding our characterization with *S. mitis* and comparing to our previously reported datasets, we continue to document repeatable and reproducible changes observed across species that can also help narrow in on critical *S. mutans* genes/operons during competitive interactions. One example is the repeated upregulation of the CRISPR2-Cas system and the downregulation of bacteriocins such as *nlmAB*, even when challenged by a high H_2_O_2_ producer. Other notable similarities included changes in the expression of genes related to carbohydrate utilization, particularly PTS genes. Recently the *S. mutans* CRISPR2-Cas system was shown to be regulated by CcpA and CodY (31), more broadly connecting the changes observed in this operon to those seen with other PTS operons, to the overall modulation of glycolytic processes for *S. mutans* in coculture. Upregulation of PTS genes and an ABC transporter operon annotated to be involved in carbohydrate transport were also upregulated in *S. mitis*, as well as *dhaL* and *dhaM*. This further supports our previous observations (16) that coculture between oral streptococci shifts metabolic preferences between species that may shift each into their own nutritional niche, a hypothesis we are further exploring.

Continued evaluation of strains undergoing competitive interactions, such as we have completed here, with further our understanding in the critical role factors like H_2_O_2_ play in active antagonism against disease-related organisms. In addition, it will continue to reveal new mechanisms such as metabolic regulation that could be utilized to gain leverage in either prebiotic or probiotic solutions as novel therapeutic strategies to prevent and/or treat dental caries. It is clear from some of our data shown here that there are several more aspects to consider when searching or selecting a potential probiotic strain for further evaluation – for example, it’s potential to disrupt pre-formed biofilms of caries pathogens and/or its impact on other commensal strains. Still, the phenotypic heterogeneity displayed between strains of the same species offers opportunities to specifically select for qualities, at the appropriate levels, that would lead to desired outcomes including the resistance to environmental perturbations or the colonization/expansion of disease-related organisms. By further surveying and charactergorizing the landscape of intermicrobial interactions, we can further our goal of utilizing natural strains and/or conditions that would lead to advances in microbiome engineering to better human health.

## MATERIALS AND METHODS

### Strain Inoculation and Growth Mediums

Overnight cultures of the bacterial strains used in this study **(Table S5)** were inoculated from single, isolated colonies on Bacto Brain Heart Infusion (Difco BHI; Fisher Bioreagents 237500) agar plates (Difco Agar, Fisher Bioreagents 214010) into BHI broth and incubated at 37°C and 5% CO_2_ with the appropriate antibiotics. Antibiotics were added to overnight growth medium BHI at 1 mg mL^-1^ for both kanamycin and spectinomycin and 0.01 mg mL^-1^ for erythromycin. The next day, cultures were harvested by centrifugation, washed to remove all traces of overnight growth medium, and normalized to OD_600 nm_ = 0.1 with 1X phosphate-buffered saline (PBS) prior to back dilution (1:100) into tryptone and yeast extract supplemented with glucose (TYG, 20 mM glucose final concentration; 10 g tryptone [Fisher Bioreagents BP1421], 5 g yeast extract [Fisher Bioreagents BP1422], 3 g K_2_HPO_4_ [Sigma-Aldrich P3786], and 3.6 g glucose per 1 L H_2_O [Sigma-Aldrich G8270]) media. 1.7 g L^-1^ sucrose (Sigma-Aldrich S7903) was added for all biofilm-related experiments (5 mM sucrose final concentration). For experiments involving catalase (Sigma-Aldrich C1345), 100 U mL^-1^ was added directly to TYG/TYGS medium prior to inoculation. For coculture competitions with *S. mutans*, strains were inoculated according to **Table S6**. Prussian blue agar plates were made as previously described (32). All strains were maintained for long-term storage at -80ºC in BHI containing 25% glycerol.

### Crystal Violet Biofilm Biomass Quantification

Strains were inoculated into 96 well plates and incubated for 24 h at 37ºC and 5% CO_2_. Following, medium from the biofilms were aspirated and plates were dunked into a bucket of water to remove all non-attached cells. After drying, 0.05 mL of 0.1% crystal violet (CV; Fisher Chemical C581) was added to each well and incubated at room temperature for 15 minutes. The CV solution were then aspirated and plates were dunked into a bucket of water again to remove excess CV. Plates were dried and imaged. Next, 0.2 mL of 30% acetic acid solution (RICCA Chemical 1383032) was added to extract the bound CV. Extracted CV solution was diluted 1:4 with water into a new 96 well plate before the absorbance at 575 nm was recorded within an Agilent Biotek Synergy H1 multimode plate reader using Gen5 microplate reader software [v 3.11 software]. All biofilm experiments were completed with at least three biological replicates measured in technical quadruplicates.

### Biofilm Microscopy

Bacterial strains were inoculated into medium that contained 1 µM Alexa Fluor 647-labeled dextran (10,000 molecular weight; anionic, fixable; Invitrogen, D22914), added to Cellvis 12 well, glass bottom, black plates (P12-1.5H-N) and incubated at 37°C and 5% CO_2_ for 24 h. Resulting biofilms were first washed with 1X PBS to remove loosely-bound cells and incubated with BSA blocking buffer at room temperature for 0.5 h (Thermo Scientific, 3% BSA is PBS; J61655.AK). Biofilms were then probed with a murine monoclonal antibody against dsDNA (Anti-dsDNA, 3519 DNA, Abcam, ab27156) [2 µg mL^-1^] within BSA blocking buffer for 1 h at room temperature. The biofilms were then washed once and incubated for 1 h at room temperature with an Alexa Fluor 594-labeled goat anti-mouse IgG highly cross-absorbed secondary antibody (Invitrogen, A32742)) [2 µg mL^-1^] within BSA blocking buffer. Finally, the biofilms were washed and stained with Hoechst 33342 solution (5 µM final concentration, Thermo Scientific, 62249) for 15 minutes (if desired). All biofilms were imaged within 1X PBS using a 40X (plan fluorite, 2.7 mm working distance, 0.6 numerical aperture) phase objective on a Agilent Biotek Lionheart FX automated microscope (Agilent Biotek, Winooski, Vermont, United States) equipped with 365 nm, 465 nm, 523 nm and 623 nm light-emitting diodes (LED; 1225007, 1225001, 1225003, 1225005) for acquiring fluorescent signals with DAPI (377/447; 1225100), GFP (469/525; 1225101), RFP (531/593; 1225103) and Cy5 (628/685; 1225105) filter cubes, respectively. Images were captured using Gen5 Image+ software, and quantification of biomass and biofilm thickness were completed either with the Gen5 Image+ software, or by importing TIF files into BiofilmQ (33). For analysis, single channel images were analyzed by setting object threshold intensity to greater than or equal to 5000 a.u. (arbitrary units) and minimum object size to greater than 5 microns. Options selected included ‘split touching objects’ and ‘fill holes in masks’. Primary edge objects were excluded from analysis. At least four images of each sample, taken at 2500 micron increments to avoid observer bias, were acquired during each experiment.

### Colony Forming Units Returned from Biofilms

1 × 10^6^ cells mL^-1^ each of *S. mutans* UA159 pMZ (kanamycin resistant) and *S. mitis* ATCC 49456 were inoculated into TYGS and incubated for 24 h at 37ºC and 5% CO_2_. After, the liquid medium was removed, representing viable cells in planktonic growth phase. The remaining attached biofilm growth phase cells were washed three times in 1X PBS to remove loosely bound cells. 1 mL of 1X PBS was added and biofilm cells were detached from 12 well polystyrene plates (Fisherbrand FB012928) using disposable cell scrapers (Fisherbrand 08100241) and transferred to a 5 mL polystyrene round-bottom tube. Cells were then sonicated within a water bath sonicator for four intervals of 30 s each while resting in between for two minutes on ice to isolate single cells. Cells were serially diluted and plated on both BHI and BHI kanamycin agar plates and incubated for 48 h at 37°C and 5% CO_2_. Colony forming units (CFUs) were later enumerated from these agar plates to determine the CFUs returned. *S. mitis* was distinguishable from *S. mutans* on BHI agar plates though colony color and morphology.

### Biofilm Formation in Spent Supernatants

Supernatants of single species or cocultures were first generated by inoculating strains in TYG medium and grown overnight, planktonically, at 37°C and 5% CO_2_. The next day, cultures were harvested by centrifugation and supernatant extracted. Supernates were treated by adjusting pH to ∼7.0 using 6N sodium hydroxide and adding back 10 mM glucose and 5 mM sucrose as a carbohydrate source. Supernatants were filtered through a 0.22 μm PES membrane filter unit (Millipore Express, SLGPR33RB) prior to inoculation of *S. mutans* to initiate biofilm growth.

### Measurement of Hydrogen Peroxide Production

Bacterial strains were first inoculated into 2 mL TYG medium in biological triplicates or quadruplicates and incubated overnight. The next morning, 0.2 mL of culture was used to measure the optical density at OD_600 nm_ with an Agilent Biotek Synergy H1 multimode plate reader using Gen5 microplate reader software [v 3.11 software]. The remaining culture was harvested by centrifugation with the supernatant saved after extraction and filtration through a 0.22 μm PES membrane filter unit (Millipore Express, SLGPR33RB). The concentration of hydrogen peroxide in the supernatants was determined using the Fluorimetric Hydrogen Peroxide Assay Kit (Sigma-Aldrich, MAK166) following the supplier’s instructions. Unknown sample values were determined using a standard curve of known hydrogen peroxide concentrations (0 – 1000 μM). Resulting concentrations were then normalized for the OD_600 nm_ of the culture at the time the supernatant was collected.

### Cloning of S. mitis ΔspxB Strain

A mutant of pyruvate oxidase (SM12261_RS06360; SM12261_1237) of *S. mitis* was created using a PCR ligation mutagenesis approach as previously described (34, 35), by replacing the open reading frame with an erythromycin antibiotic resistance cassette. The PCR ligation product (∼0.1 µg) was transformed into *S. mitis* using a 0.5 µM final concentration of synthetic CSP peptide (sCSP, amino acid sequence = EIRQTHNIFFNFFKRR, Biomatik^™^) within BHI medium and plating the culture onto BHI agar plates containing 0.01 mg/mL erythromycin. Resulting mutants were verified both via PCR as well as with whole genome sequencing (via SeqCenter). The mutant strain used in this study was verified to contain no other mutations (i.e., single nucleotide polymorphisms or SNPs) other than the target gene of interest. All primers used in the construction of the mutant strains are listed within **Table S7**.

### Addition of S. mitis to Pre-formed S. mutans Biofilms

Biofilms of *S. mutans* were incubated in TYGS medium for 24 h. At the same time, overnight cultures of *S. mitis* were inoculated. The next day, *S. mitis* cultures were harvested by centrifugation, washed in 1X PBS, and resuspended in fresh TYGS medium. The OD_600 nm_ was measured, and separate cell suspensions were adjusted to OD_600 nm_ = 1.0 (high cell density), 0.4 (medium cell density), or 0.1 (low cell density) through either further concentration of cell suspensions (i.e., for the high cell density) or dilution of cell suspensions (low cell density) using TYGS. After, the growth medium from the *S. mutans* biofilms were removed via aspiration, biofilms were washed in 1X PBS, and cell suspensions of *S. mitis* added to the biofilms prior to returning to incubation for an additional 24 h. Addition of fresh TYGS medium served as a “-*S. mitis*” control, and addition of 1X PBS to *S. mutans* biofilms served to preserve biofilm biomass that had been formed at the 24 h time point, prior to control medium or *S. mitis* addition.

### Harvesting Cultures and RNA Isolation for RNA-Seq

*S. mutans* and *S. mitis*, in either monocultures or a coculture between the two species, were grown in TYG medium until a measured optical density (OD_600 nm_) of 0.4 was reached before harvesting by centrifugation. Cell pellets were resuspended in 2 mL of RNAprotect bacterial reagent (Qiagen; 76506) and incubated at room temperature for 5 minutes prior to storage at - 80°C until further processing. For RNA isolation, the cell suspensions were thawed and RNAprotect removed by centrifugation and aspiration. Cell pellets were resuspended in a cell lysis buffer containing 0.5 mL TE buffer (Invitrogen; 12090015) along with 25 mg lysozyme (Sigma-Aldrich, L4919) and 20 units mutanolysin (Sigma-Aldrich, M9901) and incubated at 37°C for 1 h. 0.025 mL of Proteinase K solution (Qiagen; 19133) was then added to each tube and incubated at room temperature for 10 minutes. Cells suspended in lysis buffer were transferred to a 2 mL screw cap tube that contained 0.5 mL of 0.1 mm disruption beads for bacteria and 0.7 mL QIAzol™ Lysis Reagent (Qiagen; 79306). Cells were lysed via mechanical disruption using a bead beater for four rounds of 30 s each, with cells resting on ice in between each round. 0.2 mL of chloroform (Fisher Chemical, AC423550250) was added to each tube and vortexed vigorously before centrifugation at max speed for 10 minutes at 4°C. The top aqueous phase from each tube was moved into a new microcentrifuge tube (∼0.7 mL), and 0.6 mL of ice cold isopropanol was added along with 1/10^th^ volume sodium acetate solution (Invitrogen, 3 M, pH 5.2; R1181) and 1 μL GlycoBlue coprecipitant (Qiagen; AM9515). RNA was precipitated after holding overnight at -80°C and centrifugation. The RNA pellets were washed in 70% ethanol and air-dried. The resulting RNA pellets were then resuspended in RLT buffer from the RNeasy Mini Kit (Qiagen; 74524) containing 2-mercaptoethanol (Sigma-Aldrich, M3148). RNA was column purified with DNase digestion (Qiagen; 79254) according to the RNeasy Mini Kit protocol. RNA concentration was measured using a Qubit Flex Fluorometer and the Qubit RNA BR Assay Kit (Thermo Scientific; Q10210).

### RNA Sequencing and Transcriptome Analysis

RNA was sequenced through SeqCenter with their 8 M Single Reads package applied to RNA from monocultures and their 16 M Single Reads package applied to RNA from cocultures. Delivered .FASTQ files were uploaded and analyzed through Galaxy (36) with a custom pipeline (18, 37, 38) that included FASTQ Groomer [v 1.1.5], FASTQ Quality Trimmer [v 1.1.5], mapping of reads to respective genome files with Bowtie2 [v 2.3.4.3] and htseq-count [v 2.0.1] on genome features from species-specific.GFF3 files that resulted in a .CSV file containing non-normalized reads counts. All read counts were combined into a single .CSV file and uploaded to Degust (39) and edgeR analysis performed to determine log2 fold change and false discovery rates (FDR) for all genome features. The *p*-value was obtained by taking the –log10 of the FDR. Total reads for each sample and the number or percentage of reads assigned to each species is documented in **Table S8**. All genome files for this analysis were accessed through NCBI and are listed in **Table S9**.

### Graphing and Statistics

Graphing of data was completed with GraphPad Prism [version 10.1.2] software. All statistical analysis was completed within GraphPad Prism using the built-in analysis tools, including Principal Component Analysis (PCA) of RNA-Seq data, one-way or two-way ANOVA with post hoc tests (Dunnett’s or Tukey’s test) for a multiple comparison, and AUC calculations.

## Data Availability

The resulting RNA-Seq raw sequencing and data files from this study are available from NCBI-GEO (Gene Expression Omnibus) under accession number **GSE273140**.

## ACKNOLWEDGEMENTS

Several of the strains included in this manuscript were generously provided by Matthew M. Ramsey from the University of Rhode Island and by Samantha J. King from the Abigail Wexner Research Institute at Nationwide Children’s Hospital. This work was supported by a grant from the National Institute of Dental and Craniofacial Research (NIDCR) of the National Institutes of Health (NIH) R03DE031766.

## AUTHOR CONTRIBUTIONS

<u>Isabella Williams</u> – conceptualized project, collected and performed data analysis, wrote the original draft and participated in review and editing of the manuscript.

<u>Jacob Tuckerman</u> – conceptualized project, collected and performed data analysis, wrote the original draft and participated in review and editing of the manuscript.

<u>Daniel Peters</u> – collected and performed data analysis, contributed to methodology development, and participated in review and editing of the manuscript.

<u>Madisen Bangs</u> – collected and performed data analysis and participated in review and editing of the manuscript.

<u>Emily Williams</u> – collected and performed data analysis and participated in review and editing of the manuscript.

<u>Iris Shin</u> – collected and performed data analysis and participated in review and editing of the manuscript.

<u>Justin Kaspar</u> - conceptualized project, collected and performed data analysis, contributed to methodology development, wrote the original draft and participated in review and editing of the manuscript.

