## Supplemental Table 1 for "A Strain of *Streptococcus mitis* Inhibits Biofilm Formation of Caries Pathogens via Abundant Hydrogen Peroxide Production"

#### Table of Contents:

Supplemental Figures (S1-S3): Pages 2 - 4

Supplemental Tables (Table S1-S9): Pages 5 – 27

Supplemental Excel File: Contains analyzed RNA-Seq data (log2 fold change and –log10 P values) for each gene within either indicated species, used to construct Supplemental Tables 1-4.

* Co-first authors
### Corresponding author

**Mailing address:**
Division of Biosciences, The Ohio State University, College of Dentistry,
305 W. 12th Avenue, Postle Hall Rm 4185, Columbus, OH 43210.


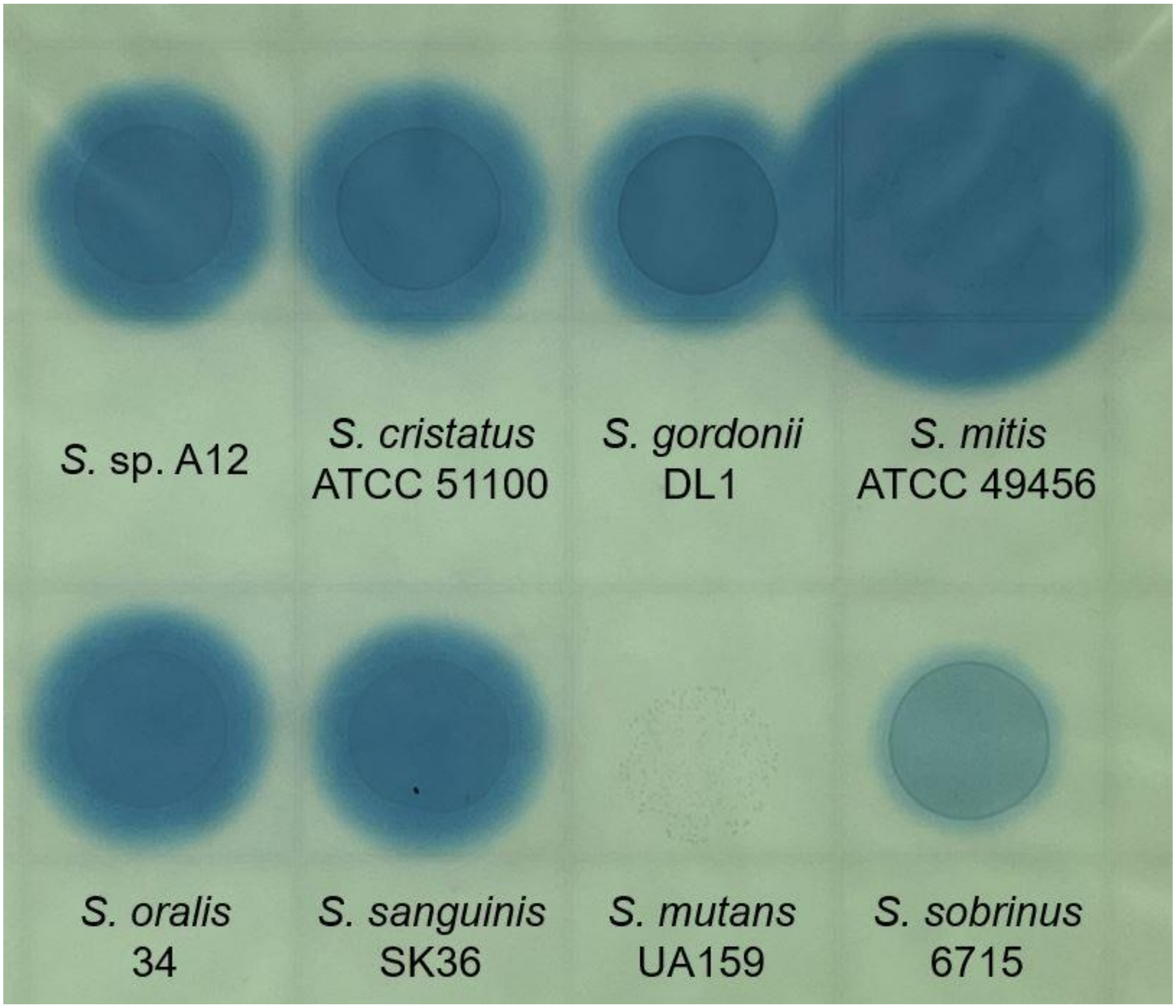

**Supplemental Figure 1. Prussian blue agar assay for hydrogen peroxide detection in different *Streptococcus* species.** Eight different species of oral streptococci were spotted (10 μL of OD600 nm = 0.1 cell suspension) on Prussian blue agar plates and incubated for 48 h at 37°C and 5% $CO_2$. The blue around each individual spotted colony represents hexacyanoferrate (III) and iron (III) present within the agar plates that precipiates in the presence of hydrogen peroxide. This experiment shows increased hydrogen peroxide production by *S. mitis* ATCC 49456 in comparison to other oral streptococci tested.

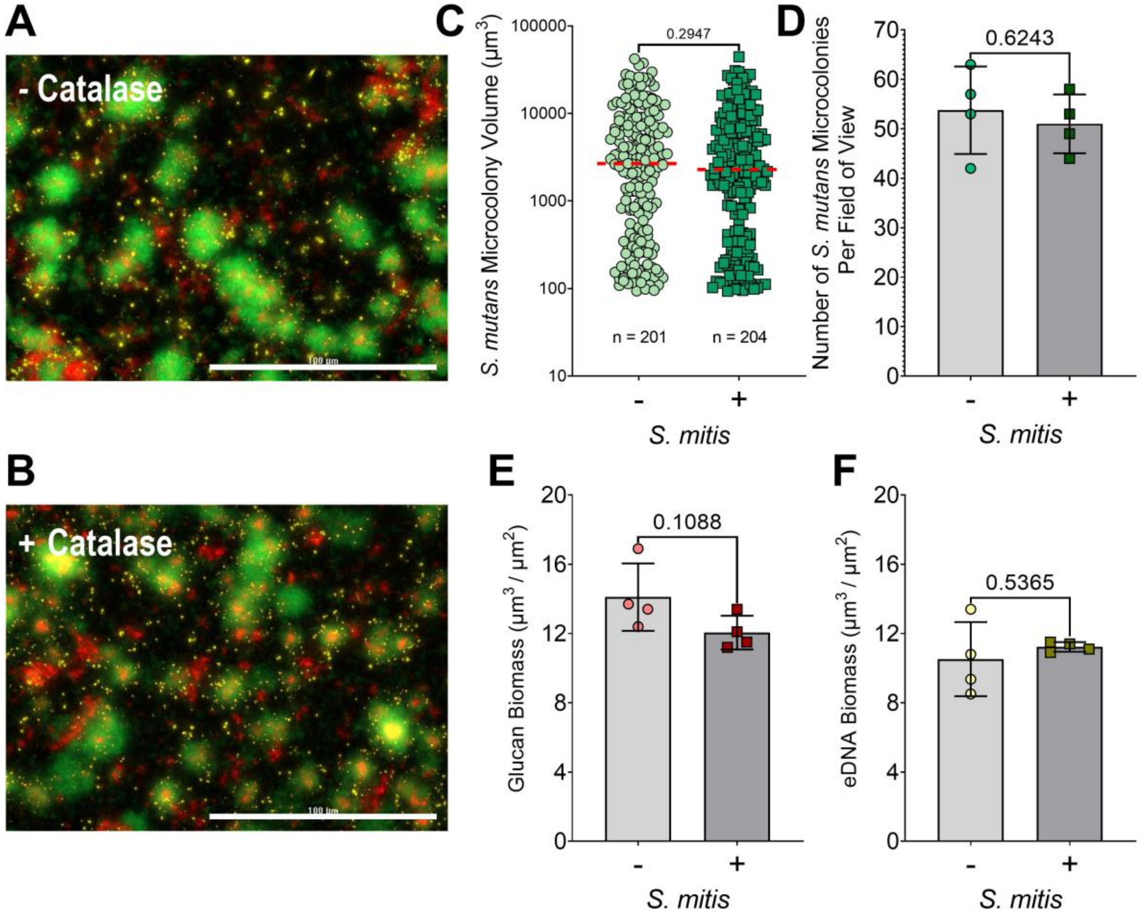

**Supplemental Figure 2. *Streptococcus mutans* monoculture biofilms in the presence and absence of 100 U mL$^{-1}$ Catalase.** **(A)** Merged representative maximum intensity 40x Z-projection of 24 h *S. mutans* monoculture biofilm grown in TYGS medium without catalase. *S. mutans* constitutively expresses GFP (green), eDNA was probed with labeled antibodies (yellow), and glucans visualized with labeled dextran (red). Scale bar (100 µm) is shown in the bottom right corner. **(B)** Merged representative maximum intensity 40X Z-projection of 24 h *S. mutans* monoculture biofilm grown in TYGS medium with addition of 100 U mL$^{-1}$ catalase. **(C)** Quantification of individual *S. mutans* microcolony volumes, **(D)** number of *S. mutans* microcolonies per field of view, **(E)** glucan biomass, and **(F)** eDNA biomass from the microscopy data shown between the two conditions. n = 4. Quantification was completed using Gen5 Image+ software. Data graphing and one-way analysis of variance (ANOVA) with multiple comparisons was completed in GraphPad Prism software. Data shows no significant differences in *S. mutans* biofilm biomass composition in the presence of catalase.

```
promoter_spxB_mitis_b6        -------tgactagttcactcttactaaactttagtaaaata---gactaaaagcgggct    50
promoter_spxB_mitis_49456     tgcaacctcaaagctgtgctttgagcaacctgtggcgagcttcctagtttgctctgtgat    60
                                     * *    *   ** * *  ** ** * *  *  *       *      * * *

promoter_spxB_mitis_b6        tgctcgcttttattataaaacagttctgtaaaacgctttcggactgtactgtaatagaga    110
promoter_spxB_mitis_49456     tttcattgagtattataaaacagaagctaaaaacgttttcaaactgtactgtaatagaga    120
                              *         *************      ****** ****  ******************

promoter_spxB_mitis_b6        atagaaacttctgagcagattcttagaaaagtcgaacgaaatgtgataaattagtagagt    170
promoter_spxB_mitis_49456     ataaaaacttctgagcagattcttagtaaagtcaaaataaatgtgataaattagtattgt    180
                              *** ********************** ****** **  ******************  **

promoter_spxB_mitis_b6        aaaaatttgctgaaacgttttcaaaaactatatgaaacttt-tttagtagatttaaaaat    229
promoter_spxB_mitis_49456     aaaaatttactgaaacgttttcaaaaactatatgaaaccttttttagtagatttaaaaat    240
                              ******** ***************************** ** ******************

promoter_spxB_mitis_b6        atttgaaggagagttatcatt    250
promoter_spxB_mitis_49456     atttgaaggagagttatcatt    261
                              *********************
```

**Supplemental Figure 3. *spxB* promoter alignment between *Streptococcus mitis* B6 and *Streptococcus mitis* ATCC 49456.** Alignment of 250 bp upstream of the *spxB* start site (ATG) in *S. mitis* B6 and 261 bp upstream in *S. mitis* ATCC 49456. The 3' end of the sequence represents the site closest to the start codon for *spxB*. Alignment was completed with the Clustal Omega multiple sequence alignment (MSA) tool with default settings.

#### SUPPLEMENTAL TABLES

**Supplemental Table 1.** Differentially expressed genes* (DEGs) between *Streptococcus mitis* ATCC 49456 grown in monoculture vs coculture with *Streptococcus mutans* UA159**.** Red, upregulated genes. Blue, downregulated genes.

*Differentially expressed genes = genes with ≥4 Log10 *P* value and Log2 fold change (FC) ≥ (-)1.

| Gene ID | Gene Name | Gene Description | Log2 Fold Change | Fold Change | P value | Log10 P value |
|---|---|---|---|---|---|---|
| SM12261_RS02695 | | sugar ABC transporter permease | 1.98 | 3.94 | 1.47E-15 | 14.8 |
| SM12261_RS02690 | | ABC transporter substrate-binding protein | 1.89 | 3.69 | 1.16E-20 | 19.9 |
| SM12261_RS09385 | | | 1.80 | 3.48 | 9.09E-09 | 8.0 |
| SM12261_RS02700 | | carbohydrate ABC transporter permease | 1.73 | 3.33 | 7.93E-11 | 10.1 |
| SM12261_RS01035 | adhE | bifunctional acetaldehyde-CoA/alcohol dehydrogenase | 1.62 | 3.08 | 4.26E-16 | 15.4 |
| SM12261_RS01275 | adhP | alcohol dehydrogenase AdhP | 1.55 | 2.94 | 1.52E-16 | 15.8 |
| SM12261_RS08975 | | ABC transporter substrate-binding protein | 1.45 | 2.74 | 2.45E-10 | 9.6 |
| SM12261_RS01820 | | CPBP family intramembrane metalloprotease | 1.40 | 2.65 | 7.63E-11 | 10.1 |
| SM12261_RS06095 | pdxT | pyridoxal 5'-phosphate synthase glutaminase subunit PdxT | 1.39 | 2.62 | 5.67E-18 | 17.2 |
| SM12261_RS02705 | | YhcH/YjgK/YiaL family protein | 1.36 | 2.57 | 7.61E-08 | 7.1 |
| SM12261_RS05425 | | sugar ABC transporter ATP-binding protein | 1.34 | 2.53 | 2.51E-10 | 9.6 |
| SM12261_RS01815 | | hypothetical protein | 1.26 | 2.39 | 1.81E-08 | 7.7 |
| SM12261_RS06100 | pdxS | pyridoxal 5'-phosphate synthase lyase subunit PdxS | 1.25 | 2.38 | 3.11E-15 | 14.5 |
| SM12261_RS07895 | | helix-turn-helix transcriptional regulator | 1.19 | 2.29 | 1.74E-07 | 6.8 |
| SM12261_RS08950 | | hypothetical protein | 1.18 | 2.27 | 6.43E-11 | 10.2 |
| SM12261_RS04745 | dhaL | dihydroxyacetone kinase subunit L | 1.18 | 2.26 | 8.49E-12 | 11.1 |
| SM12261_RS08965 | | bacteriocin immunity protein | 1.17 | 2.25 | 1.49503E-05 | 4.8 |
| SM12261_RS02685 | | PTS transporter subunit EIIC | 1.17 | 2.25 | 3.29E-10 | 9.5 |
| SM12261_RS09150 | | hypothetical protein | 1.17 | 2.24 | 2.64E-17 | 16.6 |
| SM12261_RS05415 | | substrate-binding domain-containing protein | 1.17 | 2.24 | 2.87E-10 | 9.5 |
| SM12261_RS00935 | | zinc-dependent alcohol dehydrogenase family protein | 1.16 | 2.24 | 3.09E-13 | 12.5 |
| SM12261_RS05430 | rbsD | D-ribose pyranase | 1.15 | 2.21 | 1.25752E-05 | 4.9 |
| SM12261_RS05420 | rbsC | ribose ABC transporter permease | 1.14 | 2.21 | 6.07258E-06 | 5.2 |
| SM12261_RS05225 | | glucose-1-phosphate adenylyltransferase | 1.12 | 2.18 | 2.40E-11 | 10.6 |
| SM12261_RS04750 | | PTS-dependent dihydroxyacetone kinase phosphotransferase subunit DhaM | 1.10 | 2.15 | 1.0872E-06 | 6.0 |
| SM12261_RS00630 | | sugar ABC transporter permease | 1.10 | 2.15 | 7.75E-07 | 6.1 |

| | | | | | | |
|---|---|---|---|---|---|---|
| SM12261_RS07200 | ugpC | sn-glycerol-3-phosphate ABC transporter ATP-binding protein UgpC | 1.10 | 2.14 | 2.2392E-05 | 4.6 |
| SM12261_RS08945 | | DUF4097 family beta strand repeat-containing protein | 1.03 | 2.05 | 1.95573E-06 | 5.7 |
| SM12261_RS06215 | | endonuclease/exonuclease/phosphatase family protein | 1.03 | 2.04 | 1.75E-07 | 6.8 |
| SM12261_RS09375 | | competence system sensor histidine kinase ComD | -1.09 | -2.12 | 1.97E-11 | 10.7 |
| SM12261_RS06165 | | bacteriocin immunity protein | -1.10 | -2.15 | 1.15375E-05 | 4.9 |

**Supplemental Table 2.** DEGs between *Streptococcus mutans* UA159 grown in monoculture vs coculture with *Streptococcus mitis* ATCC 49456**.** Red, upregulated genes. Blue, downregulated genes.

| Gene ID | Gene Name | Gene Description | Log2 Fold Change | Fold Change | P value | Log10 P value |
|---|---|---|---|---|---|---|
| **SMU_2027** | | transcriptional regulator/repressor | 3.70 | 13.04 | 8.73E-07 | 6.1 |
| **SMU_1470c** | | conserved hypothetical protein | 3.61 | 12.22 | 1.61E-09 | 8.8 |
| **SMU_202c** | | conserved hypothetical protein/Streptococcus-specific protein | 3.40 | 10.56 | 2.24E-06 | 5.7 |
| **SMU_206c** | | hypothetical protein | 3.25 | 9.54 | 1.31E-19 | 18.9 |
| **SMU_1888** | | transposase fragment | 3.25 | 9.50 | 6.37E-18 | 17.2 |
| **SMU_210c** | | hypothetical protein | 3.24 | 9.43 | 1.30E-14 | 13.9 |
| **SMU_1752c** | | hypothetical protein | 3.21 | 9.24 | 1.60E-14 | 13.8 |
| **SMU_207c** | | transcriptional regulator | 3.15 | 8.88 | 1.59E-16 | 15.8 |
| **SMU_1764c** | | conserved hypothetical protein | 3.14 | 8.82 | 6.55E-08 | 7.2 |
| **SMU_1762c** | | conserved hypothetical protein | 3.14 | 8.79 | 3.82E-14 | 13.4 |
| **SMU_200c** | | hypothetical protein | 3.11 | 8.60 | 6.72E-12 | 11.2 |
| **SMU_2104a** | | 50S ribosomal protein L32 | 3.08 | 8.44 | 4.04E-17 | 16.4 |
| **SMU_211c** | | hypothetical protein | 3.07 | 8.43 | 1.55E-17 | 16.8 |
| **SMU_1753c** | | conserved hypothetical protein | 3.05 | 8.26 | 2.83E-11 | 10.5 |
| **SMU_1763c** | | conserved hypothetical protein | 3.03 | 8.16 | 7.08E-15 | 14.2 |
| **SMU_714** | | translation elongation factor Tu | 3.03 | 8.15 | 8.49E-09 | 8.1 |
| **SMU_201c** | | conserved hypothetical protein | 3.01 | 8.08 | 1.77E-12 | 11.8 |
| **SMU_1754c** | | conserved hypothetical protein | 2.99 | 7.93 | 5.82E-08 | 7.2 |
| **SMU_208c** | | conserved hypothetical protein, FtsK/SpoIIIE family | 2.98 | 7.90 | 3.46E-09 | 8.5 |
| **SMU_1760c** | | conserved hypothetical protein | 2.98 | 7.87 | 7.00E-11 | 10.2 |
| **SMU_1247** | *eno* | enolase | 2.93 | 7.62 | 3.82E-06 | 5.4 |
| **SMU_1758c** | | conserved hypothetical protein | 2.87 | 7.34 | 2.54E-09 | 8.6 |
| **SMU_1755c** | | conserved hypothetical protein | 2.86 | 7.26 | 1.56E-13 | 12.8 |
| **SMU_1750c** | | hypothetical protein | 2.84 | 7.14 | 1.40E-07 | 6.9 |
| **SMU_360** | *gapC* | glyceraldehyde-3-phosphate dehydrogenase; plasmin receptor | 2.82 | 7.05 | 1.80E-08 | 7.7 |
| **SMU_2026c** | | 30S ribosomal protein S10 fragment | 2.81 | 7.01 | 2.70E-16 | 15.6 |
| **SMU_209c** | | hypothetical protein | 2.80 | 6.97 | 2.09E-11 | 10.7 |
| **SMU_1757c** | | conserved hypothetical protein | 2.76 | 6.76 | 1.74E-11 | 10.8 |
| **SMU_1761c** | | conserved hypothetical protein | 2.75 | 6.72 | 1.01E-13 | 13.0 |
| **SMU_213c** | | hypothetical protein | 2.66 | 6.33 | 1.17E-14 | 13.9 |
| **SMU_99** | *fbaA* | fructose-bisphosphate aldolase | 2.64 | 6.25 | 6.92E-15 | 14.2 |
| **SMU_205c** | | conserved hypothetical protein | 2.64 | 6.25 | 5.38E-15 | 14.3 |
| **SMU_1682c** | | conserved hypothetical protein (possible intracellular protease) | 2.61 | 6.09 | 5.82E-14 | 13.2 |
| **SMU_198c** | | conjugative transposon protein | 2.60 | 6.05 | 1.01E-10 | 10.0 |
| **SMU_215c** | | hypothetical protein | 2.50 | 5.65 | 1.56E-12 | 11.8 |
| **SMU_212c** | | hypothetical protein | 2.48 | 5.56 | 4.57E-10 | 9.3 |

| | | | | | | |
|---|---|---|---|---|---|---|
| **SMU_204c** | | hypothetical protein | 2.44 | 5.43 | 6.12E-14 | 13.2 |
| **SMU_199c** | | hypothetical protein | 2.44 | 5.41 | 1.02E-14 | 14.0 |
| **SMU_849** | | 50S ribosomal protein L27 | 2.41 | 5.33 | 1.19E-10 | 9.9 |
| **SMU_698** | | 50S ribosomal protein L35 | 2.33 | 5.01 | 6.58E-11 | 10.2 |
| **SMU_1858** | *rs18* | 30S ribosomal protein S18 | 2.27 | 4.81 | 1.60E-08 | 7.8 |
| **SMU_402** | *pfl* | pyruvate formate-lyase | 2.26 | 4.78 | 5.06E-11 | 10.3 |
| **SMU_197c** | | hypothetical protein | 2.22 | 4.66 | 2.93E-11 | 10.5 |
| **SMU_2020** | *rl16* | 50S ribosomal protein L16 | 2.21 | 4.62 | 1.47E-13 | 12.8 |
| **SMU_120** | | 50S ribosomal protein L28 | 2.20 | 4.58 | 4.55E-15 | 14.3 |
| **SMU_216c** | | hypothetical protein | 2.18 | 4.53 | 2.68E-13 | 12.6 |
| **SMU_1379** | | | 2.17 | 4.51 | 2.96E-09 | 8.5 |
| **SMU_2010** | *rl18* | 50S ribosomal protein L18 | 2.16 | 4.46 | 1.09E-13 | 13.0 |
| **SMU_961** | | macrophage infectivity potentiator-related protein | 2.13 | 4.39 | 4.14E-16 | 15.4 |
| **SMU_92c** | | | 2.11 | 4.30 | 1.77E-14 | 13.8 |
| **SMU_766** | | transposase | 2.06 | 4.17 | 4.51E-10 | 9.3 |
| **SMU_93c** | | | 2.04 | 4.12 | 1.41E-09 | 8.9 |
| **SMU_1627** | *rl11* | 50S ribosomal protein L11 | 1.98 | 3.94 | 4.90E-12 | 11.3 |
| **SMU_1781** | | conserved hypothetical protein | 1.91 | 3.76 | 6.62E-14 | 13.2 |
| **SMU_636** | | glucosamine-6-phosphate isomerase | 1.91 | 3.76 | 1.04E-06 | 6.0 |
| **SMU_962** | | acyl-CoA dehydrogenase | 1.91 | 3.75 | 5.19E-19 | 18.3 |
| **SMU_2002** | *rs11* | 30S ribosomal protein S11 | 1.88 | 3.68 | 7.09E-13 | 12.1 |
| **SMU_1782** | | conserved hypothetical protein | 1.81 | 3.50 | 1.50E-11 | 10.8 |
| **SMU_196c** | | immunogenic secreted protein (transfer protein) | 1.80 | 3.49 | 4.38E-11 | 10.4 |
| **SMU_936** | | amino acid ABC transporter, ATP-binding protein | 1.76 | 3.39 | 1.62E-05 | 4.8 |
| **SMU_94c** | | transposase fragment | 1.65 | 3.13 | 1.22E-09 | 8.9 |
| **SMU_934** | | amino acid ABC transporter, permease protein | 1.64 | 3.12 | 1.06E-13 | 13.0 |
| **SMU_865** | | 30S ribosomal protein S16 | 1.62 | 3.08 | 1.35E-19 | 18.9 |
| **SMU_1860** | *rs6* | 30S ribosomal protein S6 | 1.61 | 3.04 | 2.43E-10 | 9.6 |
| **SMU_2017** | *rl14* | 50S ribosomal protein L14 | 1.60 | 3.03 | 2.80E-15 | 14.6 |
| **SMU_1183** | *mtlA2* | phosphotransferase system enzyme II | 1.60 | 3.02 | 1.78E-16 | 15.7 |
| **SMU_753** | | conserved hypothetical protein | 1.56 | 2.95 | 1.42E-09 | 8.8 |
| **SMU_997** | | inorganic ion ABC transporter,ATP-binding protein; possible ferrichrome transport system | 1.55 | 2.93 | 1.39E-13 | 12.9 |
| **SMU_699** | | 50S ribosomal protein L20 | 1.54 | 2.92 | 6.69E-13 | 12.2 |
| **SMU_1072c** | | acyltransferase | 1.53 | 2.88 | 6.86E-11 | 10.2 |
| **SMU_589** | | histone-like DNA-binding protein | 1.53 | 2.88 | 8.79E-15 | 14.1 |
| **SMU_2021** | *rs3* | 30S ribosomal protein S3 | 1.52 | 2.88 | 1.17E-15 | 14.9 |
| **SMU_935** | | amino acid ABC transporter, permease protein | 1.51 | 2.85 | 1.33E-16 | 15.9 |
| **SMU_528c** | | conserved hypothetical protein | 1.49 | 2.82 | 9.34E-17 | 16.0 |
| **SMU_940c** | | hemolysin III-related protein | 1.49 | 2.81 | 3.88E-14 | 13.4 |
| **SMU_995** | | ferrichrome ABC transporter (permease) | 1.49 | 2.81 | 7.93E-11 | 10.1 |

| | | | | | | |
|---|---|---|---|---|---|---|
| **SMU_981** | *bglB1* | beta-glucosidase, BglB protein | 1.48 | 2.80 | 8.67E-18 | 17.1 |
| **SMU_217c** | | conserved hypothetical protein; Streptococcus-specific protein | 1.48 | 2.78 | 6.54E-12 | 11.2 |
| **SMU_933** | | amino acid ABC transporter, amino acid substrate-binding protein | 1.47 | 2.76 | 5.30E-17 | 16.3 |
| **SMU_1343c** | | polyketide synthase | 1.46 | 2.75 | 2.08E-16 | 15.7 |
| **SMU_2016** | *rl24* | 50S ribosomal protein L24 | 1.45 | 2.73 | 1.45E-17 | 16.8 |
| **SMU_1184c** | | transcriptional regulator | 1.45 | 2.72 | 1.01E-09 | 9.0 |
| **SMU_1182** | *mtlD* | mannitol 1-phosphate 5-dehydrogenase | 1.42 | 2.68 | 7.07E-17 | 16.2 |
| **SMU_1988c** | | probable DNA binding protein | 1.42 | 2.67 | 2.92E-12 | 11.5 |
| **SMU_1341c** | | gramicidin S synthase/mycosubtilin synthetase chain mycB | 1.40 | 2.64 | 7.07E-18 | 17.2 |
| **SMU_527** | | conserved hypothetical protein | 1.40 | 2.63 | 2.41E-11 | 10.6 |
| **SMU_1346** | *bacT* | thioesterase II-like protein | 1.40 | 2.63 | 4.37E-11 | 10.4 |
| **SMU_1340** | *bacA2* | bacitracin synthetase 1/ tyrocidin synthetase III | 1.39 | 2.62 | 2.11E-17 | 16.7 |
| **SMU_2025** | *rl3* | 50S ribosomal protein L3 | 1.38 | 2.60 | 7.51E-15 | 14.1 |
| **SMU_998** | | ABC transporter, ferrichrome-binding protein | 1.38 | 2.60 | 1.01E-16 | 16.0 |
| **SMU_1068c** | | ABC transporter, ATP-binding protein | 1.36 | 2.57 | 2.31E-08 | 7.6 |
| **SMU_1345c** | | peptide synthetase similar to mycA | 1.33 | 2.51 | 3.75E-10 | 9.4 |
| **SMU_2057c** | | cadmium-efflux ATPase, E1-E2 (heavy metal-transporting ATPase) | 1.32 | 2.50 | 5.03E-17 | 16.3 |
| **SMU_1335c** | | enoyl-acyl carrier protein(ACP) reductase; dioxygenase related to 2-nitropropane dioxygenase | 1.30 | 2.46 | 1.36E-15 | 14.9 |
| **SMU_1743** | *acp* | acyl carrier protein | 1.25 | 2.38 | 5.53E-13 | 12.3 |
| **SMU_1347c** | | | 1.25 | 2.38 | 6.12E-11 | 10.2 |
| **SMU_2019** | *rl29* | 50s ribosomal protein L29 | 1.24 | 2.37 | 3.51E-15 | 14.5 |
| **SMU_195c** | | hypothetical protein | 1.24 | 2.36 | 1.91E-11 | 10.7 |
| **SMU_1365c** | | | 1.22 | 2.33 | 7.84E-10 | 9.1 |
| **SMU_1342** | *bacA1* | bacitracin synthetase | 1.21 | 2.31 | 6.93E-10 | 9.2 |
| **SMU_184** | *sloC* | ABC transporter element, iron predicted binding protein | 1.20 | 2.30 | 1.07E-08 | 8.0 |
| **SMU_1339** | *bacD* | bacitracin synthetase; surfactin synthetase | 1.18 | 2.27 | 7.43E-17 | 16.1 |
| **SMU_1344c** | | malonyl CoA-acyl carrier protein transacylase | 1.18 | 2.26 | 1.31E-14 | 13.9 |
| **SMU_924** | *tpx* | thiol peroxidase | 1.17 | 2.26 | 1.66E-10 | 9.8 |
| **SMU_194c** | | conserved hypothetical protein, phage-related | 1.17 | 2.25 | 3.06E-08 | 7.5 |
| **SMU_2022** | *rl22* | 50S ribosomal protein L22 | 1.17 | 2.25 | 1.09E-18 | 18.0 |
| **SMU_193c** | | conserved hypothetical protein | 1.15 | 2.23 | 3.88E-09 | 8.4 |
| **SMU_1336** | *pksD* | conserved hypothetical protein | 1.15 | 2.23 | 4.15E-11 | 10.4 |
| **SMU_1626** | *rl1* | 50S ribosomal protein L1 | 1.15 | 2.22 | 3.50E-10 | 9.5 |
| **SMU_186** | *sloR* | metal-dependent transcriptional regulator (probable DtxR homolog) | 1.15 | 2.22 | 1.15E-15 | 14.9 |
| **SMU_925** | | bacteriocin immunity protein | 1.14 | 2.21 | 9.77E-13 | 12.0 |
| **SMU_941c** | | conserved hypothetical protein | 1.14 | 2.20 | 6.71E-13 | 12.2 |
| **SMU_185** | | hypothetical protein | 1.14 | 2.20 | 6.22E-07 | 6.2 |

| | | | | | | |
|---|---|---|---|---|---|---|
| **SMU_637c** | | hypothetical protein | 1.13 | 2.19 | 1.3704E-06 | 5.9 |
| **SMU_769** | | conserved hypothetical protein | 1.12 | 2.18 | 2.14E-11 | 10.7 |
| **SMU_996** | | ABC transporter, permease protein;possible ferrichrome transport system | 1.12 | 2.17 | 4.79E-13 | 12.3 |
| **SMU_2009** | *rs5* | 30S ribosomal protein S5 | 1.11 | 2.16 | 1.33E-16 | 15.9 |
| **SMU_540** | *dpr* | peroxide resistance protein / iron binding protein | 1.10 | 2.14 | 1.62E-14 | 13.8 |
| **SMU_2018** | *rs17* | 30S ribosomal protein S17 | 1.10 | 2.14 | 1.48E-17 | 16.8 |
| **SMU_1337c** | | alpha/beta superfamily hydrolases | 1.09 | 2.13 | 2.02E-09 | 8.7 |
| **SMU_154** | | 30S ribosomal protein S15 | 1.09 | 2.13 | 7.39E-12 | 11.1 |
| **SMU_361** | *pgk* | phosphoglycerate kinase | 1.09 | 2.13 | 1.93E-07 | 6.7 |
| **SMU_358** | | 30S ribosomal protein S7 | 1.09 | 2.13 | 7.68E-18 | 17.1 |
| **SMU_1366c** | | | 1.08 | 2.12 | 7.70E-13 | 12.1 |
| **SMU_1680c** | | hypothetical protein | 1.08 | 2.11 | 9.47E-10 | 9.0 |
| **SMU_05** | | conserved hypothetical protein | 1.04 | 2.06 | 1.40E-11 | 10.9 |
| **SMU_1641c** | | conserved hypothetical protein | 1.04 | 2.06 | 2.45E-06 | 5.6 |
| **SMU_1070c** | | conserved hypothetical protein | 1.04 | 2.06 | 3.50E-11 | 10.5 |
| **SMU_932** | | conserved hypothetical protein | 1.03 | 2.05 | 1.02E-07 | 7.0 |
| **SMU_2096c** | | conserved hypothetical protein | 1.03 | 2.05 | 5.31E-09 | 8.3 |
| **SMU_1067c** | | ABC transporter, permease protein | 1.02 | 2.03 | 1.69E-11 | 10.8 |
| **SMU_1348c** | | | 1.02 | 2.03 | 2.68E-10 | 9.6 |
| **SMU_1617** | *era* | GTP-binding protein era homolog. | 1.02 | 2.02 | 1.08E-09 | 9.0 |
| **SMU_340** | | 50S ribosomal protein L34 | 1.01 | 2.01 | 4.33E-13 | 12.4 |
| **SMU_1904c** | | hypothetical protein | -1.00 | 0.50 | 1.42E-09 | 8.8 |
| **SMU_145** | | major facilitator superfamily transporter, efflux protein | -1.00 | 0.50 | 3.33E-11 | 10.5 |
| **SMU_300** | *tgt* | tRNA-guanine transglycosylase; queuine tRNA-ribosyltransferase | -1.00 | 0.50 | 1.24E-13 | 12.9 |
| **SMU_1999c** | | glutamate--cysteine ligase | -1.00 | 0.50 | 2.56E-16 | 15.6 |
| **SMU_1390** | | conserved hypothetical protein | -1.01 | 0.50 | 1.59E-10 | 9.8 |
| **SMU_2038** | *pttB* | phosphotransferase system, trehalose-specific IIBC component (EIIBC-tre) | -1.02 | 0.49 | 5.93E-17 | 16.2 |
| **SMU_301** | | conserved hypothetical protein | -1.02 | 0.49 | 3.30E-13 | 12.5 |
| **SMU_1003** | *gid* | glucose-inhibited division protein | -1.02 | 0.49 | 7.33E-17 | 16.1 |
| **SMU_256** | *oppB* | oligopeptide ABC transporter, permease | -1.03 | 0.49 | 2.68E-10 | 9.6 |
| **SMU_1637c** | | hypothetical protein | -1.04 | 0.49 | 5.93E-11 | 10.2 |
| **SMU_532** | *trpE* | anthranilate synthase, component I | -1.04 | 0.49 | 2.65E-14 | 13.6 |
| **SMU_887** | *galT* | galactose-1-phosphate uridylyltransferase | -1.06 | 0.48 | 1.01E-09 | 9.0 |
| **SMU_892** | *hsdS* | type I restriction-modification system specificity determinant | -1.06 | 0.48 | 5.68E-14 | 13.2 |
| **SMU_1954** | *groEL* | chaperonin GroEL | -1.07 | 0.48 | 6.14E-09 | 8.2 |
| **SMU_1511c** | | acetyltransferase; possible transcriptional repressor | -1.08 | 0.47 | 6.62E-16 | 15.2 |
| **SMU_1955** | *groES* | co-chaperonin 10kDa | -1.09 | 0.47 | 2.88E-10 | 9.5 |
| **SMU_441** | | transcriptional regulator | -1.09 | 0.47 | 1.23E-08 | 7.9 |

| | | | | | | |
|---|---|---|---|---|---|---|
| **SMU_1463c** | | conserved hypothetical protein, NIF3-related | -1.09 | 0.47 | 4.69E-12 | 11.3 |
| **SMU_302** | | conserved hypothetical protein | -1.10 | 0.47 | 1.65E-12 | 11.8 |
| **SMU_915c** | | conserved hypothetical protein | -1.11 | 0.46 | 2.41E-07 | 6.6 |
| **SMU_872** | | fructose-specific PTS system enzyme IIBC component | -1.11 | 0.46 | 8.70E-11 | 10.1 |
| **SMU_378** | | hypothetical protein | -1.12 | 0.46 | 7.36979E-05 | 4.1 |
| **SMU_558** | | isoleucine-tRNA synthetase | -1.13 | 0.46 | 3.03E-15 | 14.5 |
| **SMU_886** | *galK* | galactokinase | -1.13 | 0.46 | 1.02839E-06 | 6.0 |
| **SMU_856** | *pyrR* | bifunctional protein: pyrimidine operon regulatory protein and uracil phosphoribosyltransferase | -1.14 | 0.45 | 4.07E-08 | 7.4 |
| **SMU_268** | *purA* | adenylosuccinate synthetase | -1.14 | 0.45 | 1.08E-13 | 13.0 |
| **SMU_97** | *pyrG* | CTP synthetase | -1.15 | 0.45 | 1.20E-10 | 9.9 |
| **SMU_1905c** | | hypothetical protein | -1.16 | 0.45 | 2.18207E-06 | 5.7 |
| **SMU_893** | | anticodon nuclease | -1.18 | 0.44 | 4.68E-07 | 6.3 |
| **SMU_1909c** | | hypothetical protein | -1.19 | 0.44 | 7.75E-08 | 7.1 |
| **SMU_438c** | | (R)-2-hydroxyglutaryl-CoA dehydratase activator-related protein | -1.20 | 0.44 | 3.92E-10 | 9.4 |
| **SMU_100** | | sorbose PTS system, IIB component | -1.21 | 0.43 | 2.22E-09 | 8.7 |
| **SMU_732** | | conserved hypothetical protein | -1.21 | 0.43 | 6.44E-13 | 12.2 |
| **SMU_1912c** | | hypothetical protein | -1.24 | 0.42 | 7.42E-09 | 8.1 |
| **SMU_440** | | hypothetical protein | -1.24 | 0.42 | 4.74E-11 | 10.3 |
| **SMU_957** | | 50S ribosomal protein L10 | -1.26 | 0.42 | 4.44E-12 | 11.4 |
| **SMU_153** | | hypothetical protein | -1.27 | 0.41 | 1.04E-07 | 7.0 |
| **SMU_457** | | hypothetical protein | -1.28 | 0.41 | 1.17E-07 | 6.9 |
| **SMU_1124** | *pdp* | pyrimidine-nucleoside phosphorylase | -1.28 | 0.41 | 2.21E-15 | 14.7 |
| **SMU_1595** | *cah* | carbonic anhydrase (carbonate dehydratase) | -1.29 | 0.41 | 1.70E-13 | 12.8 |
| **SMU_1907** | | hypothetical protein | -1.29 | 0.41 | 2.39E-07 | 6.6 |
| **SMU_1224** | *pyrK* | dihydroorotate dehydrogenase electron transfer subunit | -1.30 | 0.41 | 8.30E-09 | 8.1 |
| **SMU_1910c** | | hypothetical protein | -1.32 | 0.40 | 1.32E-10 | 9.9 |
| **SMU_279** | | hypothetical protein | -1.33 | 0.40 | 2.50E-08 | 7.6 |
| **SMU_1125c** | | conserved hypothetical protein | -1.34 | 0.40 | 1.19E-13 | 12.9 |
| **SMU_152** | | hypothetical protein | -1.34 | 0.39 | 5.49605E-06 | 5.3 |
| **SMU_83** | *dnaJ* | co-chaperone protein DnaJ | -1.36 | 0.39 | 8.08E-10 | 9.1 |
| **SMU_1487** | | conserved hypothetical protein | -1.37 | 0.39 | 3.93E-07 | 6.4 |
| **SMU_910** | *gtfD* | glucosyltransferase-S | -1.37 | 0.39 | 5.03E-16 | 15.3 |
| **SMU_1223** | *pyrDB* | dihydroorotate dehydrogenase | -1.38 | 0.38 | 3.38E-10 | 9.5 |
| **SMU_1554c** | | conserved hypothetical protein | -1.38 | 0.38 | 5.00062E-06 | 5.3 |
| **SMU_1913c** | | hypothetical protein; immunity protein, BLpL-like | -1.40 | 0.38 | 5.05E-10 | 9.3 |
| **SMU_2037** | *treA* | trehalose-6-phosphate hydrolase | -1.41 | 0.38 | 1.53E-11 | 10.8 |

| | | | | | | |
|---|---|---|---|---|---|---|
| **SMU_151** | | non-lantibiotic mutacin IV B | -1.42 | 0.37 | 3.98E-07 | 6.4 |
| **SMU_241c** | | amino acid ABC transporter, ATP-binding protein | -1.43 | 0.37 | 9.68E-13 | 12.0 |
| **SMU_1816c** | | transposon-related, maturase-related protein fragment | -1.45 | 0.37 | 1.23483E-06 | 5.9 |
| **SMU_859** | *pyrA* | carbamoyl-phosphate synthase,small subunit | -1.50 | 0.35 | 2.95E-10 | 9.5 |
| **SMU_150** | | non-lantibiotic mutacin IV A | -1.53 | 0.35 | 1.41E-07 | 6.8 |
| **SMU_242c** | | glutamine ABC transporter, solute binding protein | -1.53 | 0.35 | 1.33E-08 | 7.9 |
| **SMU_2146c** | | conserved hypothetical protein | -1.54 | 0.35 | 1.30E-16 | 15.9 |
| **SMU_1879** | | mannose PTS system component IID | -1.58 | 0.33 | 9.56E-09 | 8.0 |
| **SMU_496** | *cysK* | cysteine synthetase A | -1.59 | 0.33 | 5.00E-12 | 11.3 |
| **SMU_857** | | xanthine/uracil permease | -1.60 | 0.33 | 4.66E-15 | 14.3 |
| **SMU_1878** | *ptnC* | mannose PTS system component IIC | -1.60 | 0.33 | 1.04E-08 | 8.0 |
| **SMU_1877** | *ptnA* | mannose PTS system component IIAB | -1.63 | 0.32 | 2.17E-11 | 10.7 |
| **SMU_871** | *pfkB* | fructose-1-phosphate kinase | -1.63 | 0.32 | 1.85E-08 | 7.7 |
| **SMU_858** | *pyrB* | aspartate transcarbamoylase | -1.66 | 0.32 | 8.91E-14 | 13.1 |
| **SMU_1961c** | | fructose-specific Enzyme IIA component | -1.67 | 0.31 | 1.08E-08 | 8.0 |
| **SMU_1286c** | | multidrug resistance permease | -1.67 | 0.31 | 2.79E-12 | 11.6 |
| **SMU_860** | *pyrAB* | carbamoyl-phosphate synthase, large subunit | -1.76 | 0.30 | 1.80E-07 | 6.7 |
| **SMU_80** | *hrcA* | heat-inducible transcription repressor | -1.85 | 0.28 | 3.83E-15 | 14.4 |
| **SMU_1908c** | | hypothetical protein | -1.86 | 0.28 | 4.26E-11 | 10.4 |
| **SMU_1958c** | | fructose-specific Enzyme IIC component | -2.14 | 0.23 | 7.17E-08 | 7.1 |
| **SMU_81** | *grpE* | co-chaperone protein GrpE | -2.23 | 0.21 | 8.49E-09 | 8.1 |
| **SMU_870** | | lactose phosphotransferase system repressor/transcriptional repressor of the fructose operon | -2.24 | 0.21 | 3.27E-08 | 7.5 |
| **SMU_1956c** | | conserved hypothetical protein | -2.28 | 0.21 | 1.52E-06 | 5.8 |
| **SMU_1957** | | fructose-specific Enzyme IID component | -2.29 | 0.20 | 1.36E-06 | 5.9 |
| **SMU_148** | *adhE* | alcohol-acetaldehyde dehydrogenase | -2.39 | 0.19 | 1.51E-13 | 12.8 |

**Supplemental Table 3.** Distribution of upregulated DEGs (from monoculture growth) between *Streptococcus mutans* UA159 grown in quadculture (*S. mutans*, *Streptococcus gordonii* DL1, *Streptococcus oralis* 34, and *Streptococcus sanguinis* SK36; GSE209925; doi: 10.1177/00220345221145906) vs with *Streptococcus mitis* ATCC 49456 in coculture.

Red – specific to quadculture
Blue – specific to coculture w/ *S. mitis*
Purple – common between both conditions

| Gene ID | Gene Name | Gene Description | Upregulated in Quadculture? | Upregulated with *S. mitis*? |
|---|---|---|---|---|
| SMU_05 | | conserved hypothetical protein | | |
| SMU_40 | | conserved hypothetical protein | | |
| SMU_41 | | hypothetical protein (protein) | | |
| SMU_55 | | hypothetical protein | | |
| SMU_92c | | transposase fragment | | |
| SMU_93c | | transposase fragment | | |
| SMU_94c | | transposase fragment | | |
| SMU_99 | *fbaA* | fructose-bisphosphate aldolase | | |
| SMU_120 | | 50S ribosomal protein L28 | | |
| SMU_154 | | 30S ribosomal protein S15 | | |
| SMU_169 | | 50S ribosomal protein L13 | | |
| SMU_170 | | 30S ribosomal protein S9 | | |
| SMU_184 | *sloC* | ABC transporter element, iron predicted binding protein | | |
| SMU_185 | | hypothetical protein | | |
| SMU_186 | *sloR* | metal-dependent transcriptional regulator (probable DtxR homolog) | | |
| SMU_191c | | phage-related integrase | | |
| SMU_193c | | conserved hypothetical protein | | |
| SMU_194c | | conserved hypothetical protein, phage-related | | |
| SMU_195c | | hypothetical protein | | |
| SMU_196c | | immunogenic secreted protein (transfer protein) | | |
| SMU_197c | | hypothetical protein | | |
| SMU_198c | | conjugative transposon protein | | |
| SMU_199c | | hypothetical protein | | |
| SMU_200c | | hypothetical protein | | |
| SMU_201c | | conserved hypothetical protein | | |
| SMU_202c | | conserved hypothetical protein/Streptococcus-specific protein | | |
| SMU_204c | | hypothetical protein | | |

| | | |
|---|---|---|
| SMU_205c | | conserved hypothetical protein |
| SMU_206c | | hypothetical protein |
| SMU_207c | | transcriptional regulator |
| SMU_208c | | conserved hypothetical protein, FtsK/SpoIIIE family |
| SMU_209c | | hypothetical protein |
| SMU_210c | | hypothetical protein |
| SMU_211c | | hypothetical protein |
| SMU_212c | | hypothetical protein |
| SMU_213c | | hypothetical protein |
| SMU_215c | | hypothetical protein |
| SMU_216c | | hypothetical protein |
| SMU_217c | | conserved hypothetical protein; Streptococcus-specific protein |
| SMU_239c | | conserved hypothetical protein |
| SMU_241c | | amino acid ABC transporter, ATP-binding protein |
| SMU_246 | *rgpG* | polysaccharide biosynthesis protein, glycosyl transferase family 4 |
| SMU_299c | | bacteriocin peptide precursor |
| SMU_340 | | 50S ribosomal protein L34 |
| SMU_357 | | 30S ribosomal protein S12 |
| SMU_358 | | 30S ribosomal protein S7 |
| SMU_360 | *gapC* | glyceraldehyde-3-phosphate dehydrogenase; plasmin receptor |
| SMU_361 | *pgk* | phosphoglycerate kinase |
| SMU_367 | | Streptococcus-specific protein; similar to glucan-binding protein |
| SMU_402 | *pfl* | pyruvate formate-lyase |
| SMU_411c | | Streptococcus-specific protein |
| SMU_412c | | Hit-like protein |
| SMU_453 | | S-adenosyl-methyltransferase, MraW family |
| SMU_527 | | conserved hypothetical protein |
| SMU_528c | | conserved hypothetical protein |
| SMU_540 | *dpr* | peroxide resistance protein / iron binding protein |
| SMU_589 | | histone-like DNA-binding protein |
| SMU_591c | | conserved hypothetical protein |
| SMU_600c | | conserved hypothetical protein |
| SMU_635 | | conserved hypothetical protein |
| SMU_636 | | glucosamine-6-phosphate isomerase |
| SMU_637c | | hypothetical protein |
| SMU_671 | *citZ* | citrate synthase |

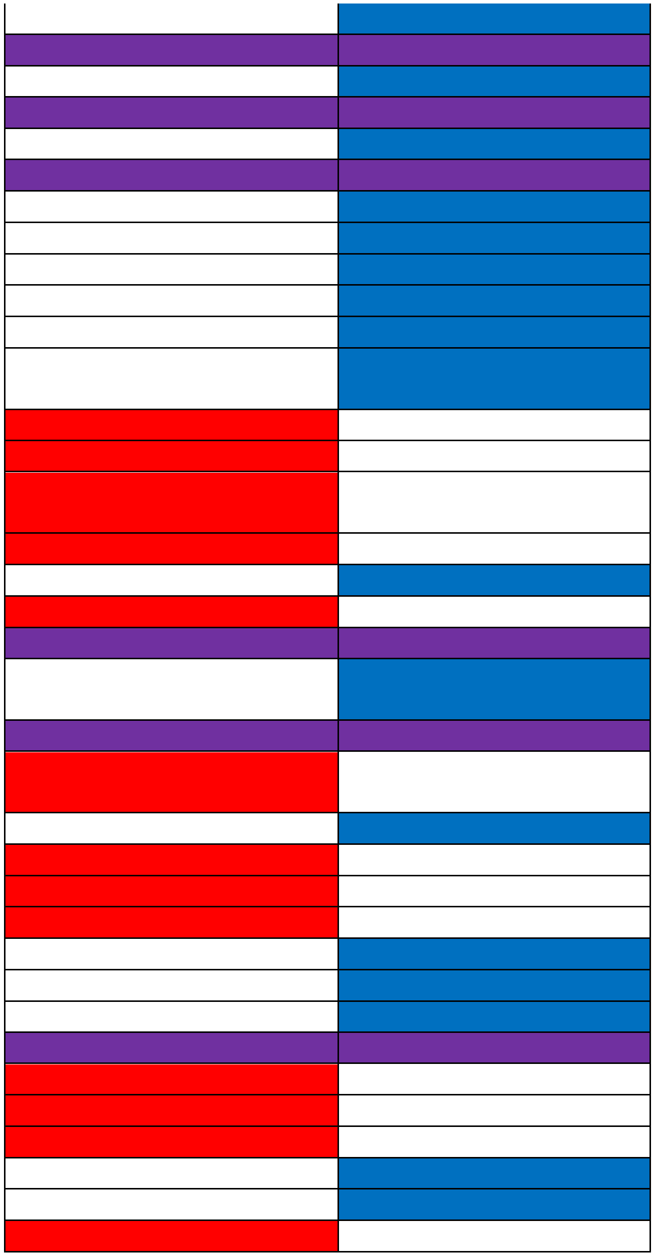

| | | |
|---|---|---|
| SMU_672 | *idh* | isocitrate dehydrogenase |
| SMU_698 | | 50S ribosomal protein L35 |
| SMU_699 | | 50S ribosomal protein L20 |
| SMU_714 | | translation elongation factor Tu |
| SMU_739c | | hypothetical protein |
| SMU_753 | | conserved hypothetical protein |
| SMU_766 | | transposase |
| SMU_769 | | conserved hypothetical protein |
| SMU_770c | | manganese transporter/possible HitA ferric iron-binding periplasmic protein |
| SMU_838 | *gshR* | glutathione reductase |
| SMU_849 | | 50S ribosomal protein L27 |
| SMU_865 | | 30S ribosomal protein S16 |
| SMU_911c | | hypothetical protein |
| SMU_924 | *tpx* | thiol peroxidase |
| SMU_925 | | bacteriocin immunity protein |
| SMU_932 | | conserved hypothetical protein |
| SMU_933 | | amino acid ABC transporter, amino acid substrate-binding protein |
| SMU_934 | | amino acid ABC transporter, permease protein |
| SMU_935 | | amino acid ABC transporter, permease protein |
| SMU_936 | | amino acid ABC transporter, ATP-binding protein |
| SMU_940c | | hemolysin III-related protein |
| SMU_941c | | conserved hypothetical protein |
| SMU_943c | | hydroxymethylglutaryl-CoA synthase |
| SMU_961 | | macrophage infectivity potentiator-related protein |
| SMU_962 | | acyl-CoA dehydrogenase |
| SMU_981 | *bglB1* | beta-glucosidase, BglB protein |
| SMU_995 | | ferrichrome ABC transporter (permease) |
| SMU_996 | | ABC transporter, permease protein;possible ferrichrome transport system |
| SMU_997 | | inorganic ion ABC transporter,ATP-binding protein; possible ferrichrome transport system |
| SMU_998 | | ABC transporter, ferrichrome-binding protein |
| SMU_1004 | *gtfB* | glucosyltransferase-I |
| SMU_1036 | | conserved hypothetical protein |
| SMU_1062 | *opuAb* | glycine-betaine binding ABC transporter permease |
| SMU_1063 | *opuAa* | amino acid ABC transporter, ATP-binding protein |
| SMU_1067c | | ABC transporter, permease protein |
| SMU_1068c | | ABC transporter, ATP-binding protein |

| | | |
|---|---|---|
| SMU_1070c | | conserved hypothetical protein |
| SMU_1072c | | acyltransferase |
| SMU_1091 | *wapE* | hypothetical protein (possible cell wall protein) |
| SMU_1131c | | hypothetical protein |
| SMU_1173 | *cysD* | O-acetylhomoserine sulfhydrylase |
| SMU_1175 | | sodium:alanine (or glycine) symporter |
| SMU_1182 | *mtlD* | mannitol 1-phosphate 5-dehydrogenase |
| SMU_1183 | *mtlA2* | phosphotransferase system enzyme II |
| SMU_1184c | | transcriptional regulator |
| SMU_1189c | | conserved hypothetical protein |
| SMU_1197 | | conserved hypothetical protein |
| SMU_1247 | *eno* | enolase |
| SMU_1322 | *budC* | acetoin reductase |
| SMU_1335c | | enoyl-acyl carrier protein(ACP) reductase; dioxygenase related to 2-nitropropane dioxygenase |
| SMU_1336 | *pksD* | conserved hypothetical protein |
| SMU_1337c | | alpha/beta superfamily hydrolases |
| SMU_1339 | *bacD* | bacitracin synthetase; surfactin synthetase |
| SMU_1340 | *bacA2* | bacitracin synthetase 1/ tyrocidin synthetase III |
| SMU_1341c | | gramicidin S synthase/mycosubtilin synthetase chain mycB |
| SMU_1342 | *bacA1* | bacitracin synthetase |
| SMU_1343c | | polyketide synthase |
| SMU_1344c | | malonyl CoA-acyl carrier protein transacylase |
| SMU_1345c | | peptide synthetase similar to mycA |
| SMU_1346 | *bacT* | thioesterase II-like protein |
| SMU_1347c | | permease |
| SMU_1348c | | ABC transporter |
| SMU_1365c | | permease |
| SMU_1366c | | ABC transporter |
| SMU_1379 | | hypothetical protein |
| SMU_1395c | | hypothetical protein |
| SMU_1396 | *gbpC* | glucan-binding protein C |
| SMU_1467 | *apt* | adenine phosphoribosyltransferase |
| SMU_1470c | | conserved hypothetical protein |
| SMU_1587c | | hypothetical protein |
| SMU_1602 | | NAD(P)H-flavin oxidoreductase |
| SMU_1603 | *lguL* | lactoylglutathione lyase |
| SMU_1617 | *era* | GTP-binding protein era homolog. |
| SMU_1626 | *rl1* | 50S ribosomal protein L1 |

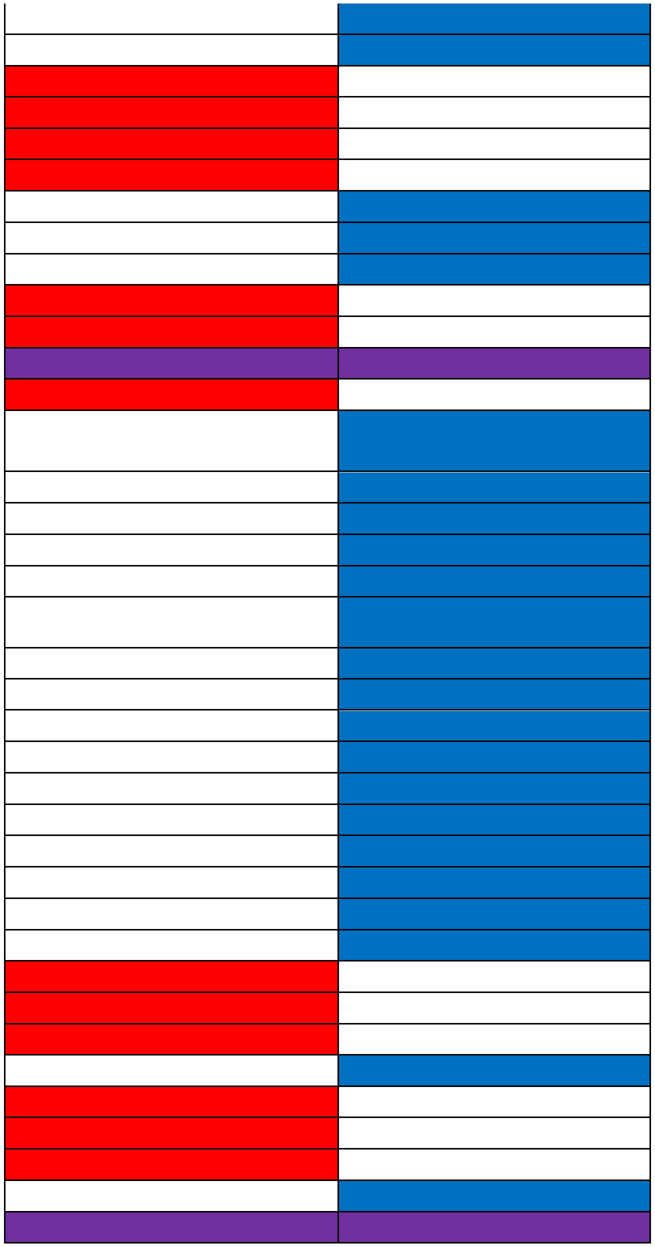

| | | |
|---|---|---|
| SMU_1627 | *rl11* | 50S ribosomal protein L11 |
| SMU_1641c | | conserved hypothetical protein |
| SMU_1649 | *exoA* | exodeoxyribonuclease III/ Smx nuclease |
| SMU_1657c | | nitrogen regulatory protein PII |
| SMU_1658 | *nrgA* | ammonium transporter, NrgA protein |
| SMU_1680c | | hypothetical protein |
| SMU_1682c | | conserved hypothetical protein (possible intracellular protease) |
| SMU_1743 | *acp* | acyl carrier protein |
| SMU_1750c | | hypothetical protein |
| SMU_1752c | | hypothetical protein |
| SMU_1753c | | conserved hypothetical protein |
| SMU_1754c | | conserved hypothetical protein |
| SMU_1755c | | conserved hypothetical protein |
| SMU_1757c | | conserved hypothetical protein |
| SMU_1758c | | conserved hypothetical protein |
| SMU_1760c | | conserved hypothetical protein |
| SMU_1761c | | conserved hypothetical protein |
| SMU_1762c | | conserved hypothetical protein |
| SMU_1763c | | conserved hypothetical protein |
| SMU_1764c | | conserved hypothetical protein |
| SMU_1781 | | conserved hypothetical protein |
| SMU_1782 | | conserved hypothetical protein |
| SMU_1802c | | conserved hypothetical protein |
| SMU_1858 | *rs18* | 30S ribosomal protein S18 |
| SMU_1860 | *rs6* | 30S ribosomal protein S6 |
| SMU_1862 | | hypothetical protein |
| SMU_1888 | | transposase fragment |
| SMU_1889c | | hypothetical protein (possible relation to bacteriocin BlpU) |
| SMU_1895c | | hypothetical protein |
| SMU_1896c | | hypothetical protein |
| SMU_1897 | | ABC transporter, ATP-binding protein; similar to BlpA |
| SMU_1898 | | ABC transporter, ATP-binding and permease element |
| SMU_1915 | *comC* | S. mutans specific competence stimulating peptide, precursor |
| SMU_1945 | | conserved hypothetical protein |
| SMU_1946 | | conserved hypothetical protein |
| SMU_1988c | | probable DNA binding protein |
| SMU_2002 | *rs11* | 30S ribosomal protein S11 |

| | | |
|---|---|---|
| SMU_2009 | *rs5* | 30S ribosomal protein S5 |
| SMU_2010 | *rl18* | 50S ribosomal protein L18 |
| SMU_2016 | *rl24* | 50S ribosomal protein L24 |
| SMU_2017 | *rl14* | 50S ribosomal protein L14 |
| SMU_2018 | *rs17* | 30S ribosomal protein S17 |
| SMU_2019 | *rl29* | 50s ribosomal protein L29 |
| SMU_2020 | *rl16* | 50S ribosomal protein L16 |
| SMU_2021 | *rs3* | 30S ribosomal protein S3 |
| SMU_2022 | *rl22* | 50S ribosomal protein L22 |
| SMU_2023c | | 30S ribosomal protein S19, C-terminal fragment |
| SMU_2025 | *rl3* | 50S ribosomal protein L3 |
| SMU_2026c | | 30S ribosomal protein S10 fragment |
| SMU_2027 | | transcriptional regulator/repressor |
| SMU_2028 | *sacB* | fructosyltransferase |
| SMU_2057c | | cadmium-efflux ATPase, E1-E2 (heavy metal-transporting ATPase) |
| SMU_2093 | *argR* | arginine repressor |
| SMU_2096c | | conserved hypothetical protein |
| SMU_2104a | | 50S ribosomal protein L32 |
| SMU_2116 | *opuCa* | glycine betaine / carnitine / choline ABC transporter, ATP-binding protein, opuCA |
| SMU_2117 | *opuCb* | glycine betaine / carnitine / choline ABC transporter permease |
| SMU_2118 | *opuCc* | glycine betaine/carnitine/choline ABC transporter, substrate-binding protein |
| SMU_2119 | *opuCd* | ABC transport betaine/carnitine/choline permease |
| SMU_2146c | | conserved hypothetical protein |
| SMU_2147c | | conserved hypothetical protein |

**Supplemental Table 4.** Distribution of downregulated DEGs (from monoculture growth) between *Streptococcus mutans* UA159 grown in quadculture (*S. mutans*, *Streptococcus gordonii* DL1, *Streptococcus oralis* 34, and *Streptococcus sanguinis* SK36; GSE209925; doi: 10.1177/00220345221145906) vs with *Streptococcus mitis* ATCC 49456 in coculture.

Red – specific to quadculture
Blue – specific to coculture w/ *S. mitis*
Purple – common between both conditions

| Gene ID | Gene Name | Gene Description | Downregulated in Quadculture? | Downregulated with *S. mitis*? |
|---|---|---|---|---|
| SMU_21 | *mreD* | cell shape-determining protein MreD | | |
| SMU_29 | | phosphoribosylaminoimidazole-succinocarboxamide synthase | | |
| SMU_78 | *fruA* | fructan hydrolase; exo-beta-D-fructosidase | | |
| SMU_80 | *hrcA* | heat-inducible transcription repressor | | |
| SMU_81 | *grpE* | co-chaperone protein GrpE | | |
| SMU_83 | *dnaJ* | co-chaperone protein DnaJ | | |
| SMU_97 | *pyrG* | CTP synthetase | | |
| SMU_100 | | sorbose PTS system, IIB component | | |
| SMU_145 | | major facilitator superfamily transporter, efflux protein | | |
| SMU_148 | *adhE* | alcohol-acetaldehyde dehydrogenase | | |
| SMU_150 | | non-lantibiotic mutacin IV A | | |
| SMU_151 | | non-lantibiotic mutacin IV B | | |
| SMU_152 | | hypothetical protein | | |
| SMU_153 | | hypothetical protein | | |
| SMU_231 | *ilvB* | acetolactate synthase, large subunit (AHAS) | | |
| SMU_233 | *ilvC* | ketol-acid reductoisomerase | | |
| SMU_241c | | amino acid ABC transporter, ATP-binding protein | | |
| SMU_242c | | glutamine ABC transporter, solute binding protein | | |
| SMU_256 | *oppB* | oligopeptide ABC transporter, permease | | |
| SMU_268 | *purA* | adenylosuccinate synthetase | | |
| SMU_279 | | hypothetical protein | | |
| SMU_300 | *tgt* | tRNA-guanine transglycosylase; queuine tRNA-ribosyltransferase | | |
| SMU_301 | | conserved hypothetical protein | | |
| SMU_302 | | conserved hypothetical protein | | |
| SMU_378 | | hypothetical protein | | |
| SMU_419 | | conserved hypothetical protein | | |
| SMU_423 | | possible bacteriocin | | |

| | | |
|---|---|---|
| SMU_438c | | (R)-2-hydroxyglutaryl-CoA dehydratase activator-related protein |
| SMU_440 | | hypothetical protein |
| SMU_441 | | transcriptional regulator |
| SMU_454 | *ftsL* | cell division protein |
| SMU_457 | | hypothetical protein |
| SMU_496 | *cysK* | cysteine synthetase A |
| SMU_525 | | ABC transporter ATP-binding / permease protein |
| SMU_532 | *trpE* | anthranilate synthase, component I |
| SMU_558 | | isoleucine-tRNA synthetase |
| SMU_574c | | effector of murein hydrolase |
| SMU_575c | | murein hydrolase regulator |
| SMU_609 | | cell wall protein precursor |
| SMU_691 | *pepT* | peptidase T (tripeptidase) |
| SMU_732 | | conserved hypothetical protein |
| SMU_856 | *pyrR* | bifunctional protein: pyrimidine operon regulatory protein and uracil phosphoribosyltransferase |
| SMU_857 | | xanthine/uracil permease |
| SMU_858 | *pyrB* | aspartate transcarbamoylase |
| SMU_859 | *pyrA* | carbamoyl-phosphate synthase,small subunit |
| SMU_860 | *pyrAB* | carbamoyl-phosphate synthase, large subunit |
| SMU_867 | *rimM* | 16S rRNA processing protein |
| SMU_870 | | lactose phosphotransferase system repressor/transcriptional repressor of the fructose operon |
| SMU_871 | *pfkB* | fructose-1-phosphate kinase |
| SMU_872 | | fructose-specific PTS system enzyme IIBC component |
| SMU_886 | *galK* | galactokinase |
| SMU_887 | *galT* | galactose-1-phosphate uridylyltransferase |
| SMU_892 | *hsdS* | type I restriction-modification system specificity determinant |
| SMU_893 | | anticodon nuclease |
| SMU_910 | *gtfD* | glucosyltransferase-S |
| SMU_915c | | conserved hypothetical protein |
| SMU_930c | | transcriptional regulator |
| SMU_940c | | hemolysin III-related protein |
| SMU_941c | | conserved hypothetical protein |
| SMU_957 | | 50S ribosomal protein L10 |
| SMU_1003 | *gid* | glucose-inhibited division protein |
| SMU_1124 | *pdp* | pyrimidine-nucleoside phosphorylase |

| | | |
|---|---|---|
| SMU_1125c | | conserved hypothetical protein |
| SMU_1134c | | phosphate ABC transporter, ATP-binding protein |
| SMU_1177c | | amino acid ABC transporter, amino acid-binding protein |
| SMU_1178c | | amino acid ABC transporter, ATP-binding protein |
| SMU_1223 | *pyrDB* | dihydroorotate dehydrogenase |
| SMU_1224 | *pyrK* | dihydroorotate dehydrogenase electron transfer subunit |
| SMU_1237c | | hypothetical protein |
| SMU_1246c | | transcriptional regulator (TetR/AcrR family) |
| SMU_1254 | | phosphatase |
| SMU_1271 | *hisG* | ATP phosphoribosyltransferase |
| SMU_1286c | | multidrug resistance permease |
| SMU_1334 | *sfp* | phosphopantetheinyl transferase |
| SMU_1335c | | enoyl-acyl carrier protein(ACP) reductase; dioxygenase related to 2-nitropropane dioxygenase |
| SMU_1337c | | alpha/beta superfamily hydrolases |
| SMU_1338c | | ABC transport macrolide permease |
| SMU_1339 | *bacD* | bacitracin synthetase; surfactin synthetase |
| SMU_1340 | *bacA2* | bacitracin synthetase 1/ tyrocidin synthetase III |
| SMU_1341c | | gramicidin S synthase/mycosubtilin synthetase chain mycB |
| SMU_1342 | *bacA1* | bacitracin synthetase |
| SMU_1343c | | polyketide synthase |
| SMU_1344c | | malonyl CoA-acyl carrier protein transacylase |
| SMU_1345c | | peptide synthetase similar to mycA |
| SMU_1346 | *bacT* | thioesterase II-like protein |
| SMU_1348c | | ABC transporter |
| SMU_1367c | | conserved hypothetical protein |
| SMU_1390 | | conserved hypothetical protein |
| SMU_1402c | | conserved hypothetical protein |
| SMU_1404c | | conserved hypothetical protein |
| SMU_1435c | | hypothetical protein |
| SMU_1463c | | conserved hypothetical protein, NIF3-related |
| SMU_1464c | | conserved hypothetical protein |
| SMU_1487 | | conserved hypothetical protein |
| SMU_1511c | | acetyltransferase; possible transcriptional repressor |
| SMU_1532 | *atpF* | ATPase, b subunit |
| SMU_1554c | | conserved hypothetical protein |
| SMU_1595 | *cah* | carbonic anhydrase (carbonate dehydratase) |
| SMU_1615c | | conserved hypothetical protein |
| SMU_1637c | | hypothetical protein |

| | | |
|---|---|---|
| SMU_1732c | | conserved hypothetical protein |
| SMU_1734 | *accA* | acetyl-CoA carboxylase alpha subunit |
| SMU_1735 | *accD* | acetyl-CoA carboxylase beta subunit |
| SMU_1816c | | transposon-related, maturase-related protein fragment |
| SMU_1877 | *ptnA* | mannose PTS system component IIAB |
| SMU_1878 | *ptnC* | mannose PTS system component IIC |
| SMU_1879 | | mannose PTS system component IID |
| SMU_1903c | | hypothetical protein |
| SMU_1904c | | hypothetical protein |
| SMU_1905c | | hypothetical protein |
| SMU_1906c | | bacteriocin-related protein |
| SMU_1907 | | hypothetical protein |
| SMU_1908c | | hypothetical protein |
| SMU_1909c | | hypothetical protein |
| SMU_1910c | | hypothetical protein |
| SMU_1912c | | hypothetical protein |
| SMU_1913c | | hypothetical protein; immunity protein, BLpL-like |
| SMU_1914c | | bacteriocin protein, BlpO-like |
| SMU_1941 | *atmB* | ABC transporter solute-binding protein |
| SMU_1954 | *groEL* | chaperonin GroEL |
| SMU_1955 | *groES* | co-chaperonin 10kDa |
| SMU_1956c | | conserved hypothetical protein |
| SMU_1957 | | fructose-specific Enzyme IID component |
| SMU_1958c | | fructose-specific Enzyme IIC component |
| SMU_1960c | | fructose-specific Enzyme IIB component |
| SMU_1961c | | fructose-specific Enzyme IIA component |
| SMU_1999c | | glutamate--cysteine ligase |
| SMU_2019 | *rl29* | 50s ribosomal protein L29 |
| SMU_2020 | *rl16* | 50S ribosomal protein L16 |
| SMU_2037 | *treA* | trehalose-6-phosphate hydrolase |
| SMU_2038 | *pttB* | phosphotransferase system, trehalose-specific IIBC component (EIIBC-tre) |
| SMU_2053c | | hypothetical protein |
| SMU_2146c | | conserved hypothetical protein |

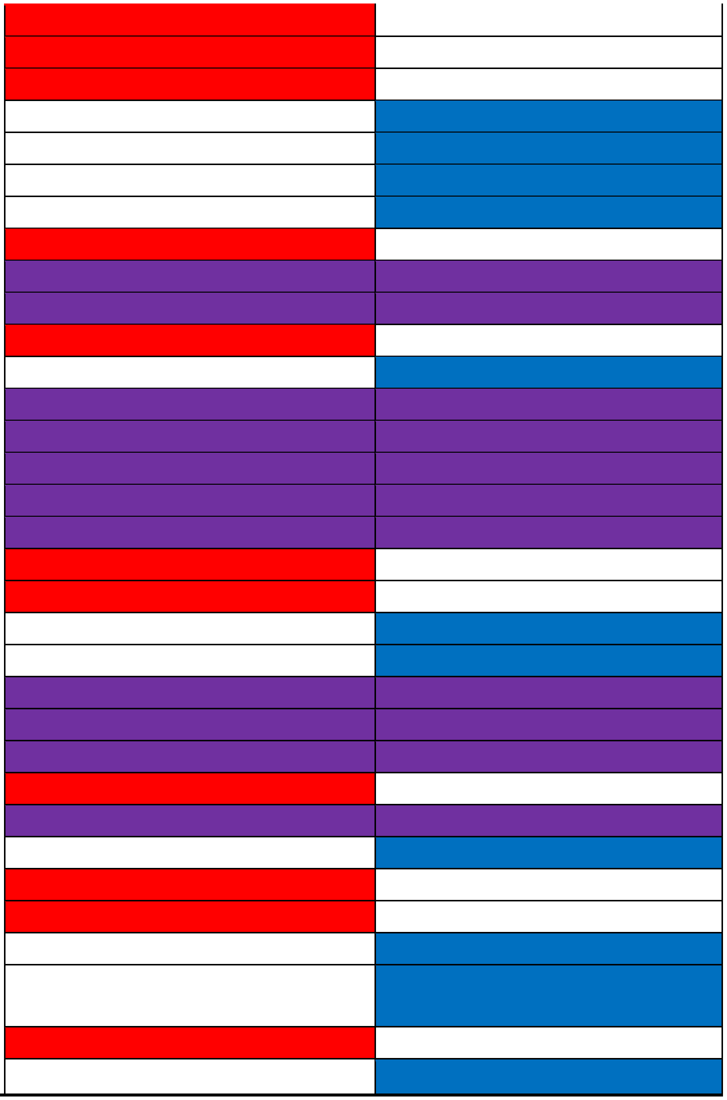

**Supplemental Table 5.** Bacterial strains used in this study.

| Species | Strain | Genotype or description | Antibiotic resistance | Reference or source |
|---|---|---|---|---|
| *Actinomyces oris* | WVU627 | | | Lab Stock |
| *Streptococcus sp.* | A12 | | | Lab Stock |
| *Streptococcus cristatus* | ATCC 51100 | | | Lab Stock |
| *Streptococcus gordonii* | DL1 | | | Lab Stock |
| *Streptococcus mits* | ATCC 49456 | | | Lab Stock |
| | 49456 Δ*spxB* | Allelic exchange of pyruvate oxidase gene (SM12261_RS06360, SM12261_1237) with an antibiotic resistant cassette | Erythromycin | This manuscript |
| | B6 | | | Lab Stock |
| | SK306 | | | Lab Stock |
| | SK569 | | | Lab Stock |
| *Streptococcus mutans* | UA159 (SMU159) | | | Lab stock |
| | 1ID3 (SMU009) | | | Lab stock |
| | 15JP3 (SMU020) | | | Lab stock |
| | 2VS1 (SMU041) | | | Lab stock |
| | NVAB (SMU053) | | | Lab stock |
| | NMT4863 (SMU057) | | | Lab stock |
| | A19 (SMU058) | | | Lab stock |
| | N66 (SMU076) | | | Lab stock |
| | SF1 (SMU080) | | | Lab stock |
| | SM6 (SMU082) | | | Lab stock |
| | U2A (SMU086) | | | Lab stock |
| | NLML1 (SMU089) | | | Lab stock |
| | 21 (SMU093) | | | Lab stock |
| | SM1 (SMU098) | | | Lab stock |
| | U2B (SMU101) | | | Lab stock |
| | SA41 (SMU104) | | | Lab stock |
| | SF12 (SMU105) | | | Lab stock |
| | R221 (SMU107) | | | Lab stock |
| | M230 (SMU108) | | | Lab stock |
| | OMZ175 (SMU109) | | | Lab stock |
| | pMZ- / UA159 | UA159 with pMZ plasmid integrated into genome. Serves as a "marked" *S. mutans* strain during colony forming unit (CFU) assays | Kanamycin | Sheilds et al. 2019, Appl Environ Microbiol. |
| | pMZ-P*veg*::*gfp* / UA159 | UA159 with pMZ plasmid integrated into genome harboring constitutively-active *gfp*. Serves as GFP(+) strain used in coculture microscopy experiments | Kanamycin | Sheilds et al. 2019, Appl Environ Microbiol. |
| *Streptococcus oralis* | 34 | | | Lab Stock |
| *Streptococcus sanguinis* | SK36 | | | Lab Stock |
| *Streptococcus sobrinus* | 6715 | | | Lab Stock |

**Supplemental Table 6.** Concentration* of inoculated bacterial strains during competitive competitions.

* Concentrations were determined based on the growth rate of individual species, such that the initial growth rate and entrance into exponential growth phase is matched to that of *Streptococcus mutans* pMZ-P*veg*::*gfp* / UA159 strain.

from doi: 10.1128/msphere.00771-23

| Strain Name | Concentration of Competitor | *S. mutans* / *S. mitis* Concentration |
|---|---|---|
| *S.* sp. A12 | 0.5x | 1x |
| *S. cristatus* ATCC 51100 | 10x | 1x |
| *S. gordonii* DL1 | 10% solution (1:10 dilution); 1x | 1x |
| *S. mitis* ATCC 49456 | 1x | 1x |
| *S. oralis* 34 | 1x | 1x |
| *S. sanguinis* SK36 | 1x | 1x |
| *S. sobrinus* 6715 | 2x | 1x |

**Supplemental Table 7.** Primers used in this study.

***Bold and underline** denotes BamHI cut site used in the PCR ligation mutagenesis approach

| Primer Name | Primer Sequence (5' - 3') * | Tm |
|---|---|---|
| mitis-spxB-A | TGC CAT TGC CGA TCA GAT CGG | 60.1 |
| mitis-spxB-B | CAT **GGA TCC** TTG CTG CAG ATG CAG | 60.4 |
| mitis-spxB-C | CGA **GGA TCC** AGA ACT CGT ACC ATT CCG | 61.8 |
| mitis-spxB-D | GTA ATC GGG ATC ATT GCT GGT AGT GC | 59.5 |

**Supplemental Table 8.** RNA-Seq read counts assigned to each species. The first table expresses total number of raw read counts for each species per library, wile the second table is expressed in % of counts within the library.

| Sample | Sample Name | *S. mutans* Counts | *S. mitis* Counts | Unassigned Counts | Total Counts |
|---|---|---|---|---|---|
| 1 | SMU_Mono_TYG_1 | 28,039,445 | - | 197,509 | **28,236,954** |
| 2 | SMU_Mono_TYG_2 | 26,483,264 | - | 203,762 | **26,687,026** |
| 3 | SMU_Mono_TYG_3 | 24,111,846 | - | 200,892 | **24,312,738** |
| 4 | Mitis_Mono_TYG_1 | - | 26,330,498 | 234,772 | **26,565,270** |
| 5 | Mitis_Mono_TYG_2 | - | 22,977,370 | 148,578 | **23,125,948** |
| 6 | Mitis_Mono_TYG_3 | - | 25,646,326 | 146,406 | **25,792,732** |
| 7 | SMU_CoMitis_TYG_1 | 16,051,834 | 30,664,536 | 98,648 | **45,632,834** |
| 8 | SMU_CoMitis_TYG_2 | 15,442,671 | 31,052,053 | 97,537 | **45,571,812** |
| 9 | SMU_CoMitis_TYG_3 | 15,502,895 | 29,075,512 | 94,681 | **43,596,136** |
| **Sample** | **Sample Name** | **SMU % Counts** | **So34 % Counts** | **Unassigned % Counts** | **Total % Counts** |
| 1 | SMU_Mono_TYG_1 | 99.3% | 0.0% | 0.7% | 100.0% |
| 2 | SMU_Mono_TYG_2 | 99.2% | 0.0% | 0.8% | 100.0% |
| 3 | SMU_Mono_TYG_3 | 99.2% | 0.0% | 0.8% | 100.0% |
| 4 | Mitis_Mono_TYG_1 | 0.0% | 99.1% | 0.9% | 100.0% |
| 5 | Mitis_Mono_TYG_2 | 0.0% | 99.4% | 0.6% | 100.0% |
| 6 | Mitis_Mono_TYG_3 | 0.0% | 99.4% | 0.6% | 100.0% |
| 7 | SMU_CoMitis_TYG_1 | 35.2% | 67.2% | 0.2% | 102.6% |
| 8 | SMU_CoMitis_TYG_2 | 33.9% | 68.1% | 0.2% | 102.2% |
| 9 | SMU_CoMitis_TYG_3 | 35.6% | 66.7% | 0.2% | 102.5% |

**Supplemental Table 9.** Genome files used in this study.

| Species | Isolate | Accession | Reference? | Source |
|---|---|---|---|---|
| *Streptococcus mutans* | UA159 | NC_004350.2 | Yes | NCBI GenBank |
| *Streptococcus mitis* | NCTC 12261 (ATCC 49456) | CP028414.1 | No | NCBI GenBank |
